# Proteomics Dissection of Cardiac Protein Profiles of Humans and Model Organisms

**DOI:** 10.1101/2020.01.08.897595

**Authors:** Nora Linscheid, Alberto Santos, Pi Camilla Poulsen, Robert W. Mills, Christian Stolte, Ulrike Leurs, Johan Z. Ye, Kirstine Calloe, Morten B. Thomsen, Bo H. Bentzen, Pia R. Lundegaard, Morten S. Olesen, Lars J. Jensen, Jesper V. Olsen, Alicia Lundby

**Affiliations:** Department of Biomedical Sciences, Faculty of Health and Medical Sciences, University of Copenhagen; The Novo Nordisk Foundation Center for Protein Research, Faculty of Health and Medical Sciences, University of Copenhagen; Department of Veterinary Science, Faculty of Health and Medical Sciences, University of Copenhagen; New York Genome Center, 101 Avenue of the Americas, New York, NY 1013, USA

## Abstract

The study of human cardiac pathologies often relies on research conducted in model organisms to gain molecular insight into disease and to develop novel treatment strategies; however, translating findings from model organisms back to human can present a significant challenge, in part due to a lack of knowledge about the differences across species in cardiac protein abundances and their interactions. Here we set out to bridge this knowledge gap by presenting a global analysis of cardiac protein expression profiles in humans and commonly used model organisms. Using quantitative mass spectrometry-based proteomics, we measured the abundance of ~7,000 proteins in samples from the separate chambers of human, pig, horse, rat, mouse and zebrafish hearts. This knowledgebase of cardiac protein signatures is accessible through an online database at: atlas.cardiacproteomics.com. Quantitative comparison of the protein profiles support the pig as model organism of choice for arrhythmogenic right ventricular cardiomyopathy whereas comparison of profiles from the two-chambered zebrafish heart suggests a better resemblance to the right side of mammalian hearts. This proteomics resource facilitates translational prospect of cardiac studies from model organisms to humans by enabling direct comparison of disease-linked protein networks across species.

## Introduction

Cardiovascular disease is the leading cause of death in the Western world, driving a pressing need to improve diagnostic and treatment options. However, limited availability of both healthy and diseased human tissue samples presents a significant challenge for studies of cardiac diseases. Even in cases where human tissue material can be obtained, complex patient phenotypes reduce experimental reproducibility and restrict the utility of those tissues. Hence, experimental studies in model systems are key in investigating the pathophysiological mechanisms underlying cardiac disease and in aiding development of diagnostics and therapeutic biomarker discovery^1,2^.

The presumption underlying any cardiac disease study conducted in a model organism is that the model adequately recapitulates relevant human cardiac physiology. Yet, macroscopic differences such as heart size and heart rate are readily apparent and corresponding differences in molecular architecture can be anticipated. Disease models have been developed in diverse organisms as each leverages some inherent advantage, for instance macrostructural similarity to human or genetic tractability, and selection of a model organism for a particular study must consider the ease of use and resources required to perform experimental manipulations and measurements as well as historical precedence. Pigs and dogs possess a high degree of similarity with human heart (electro)physiology and accordingly have been used extensively for modeling cardiac disease as well as for testing therapeutic treatments^3^. In general, large mammals are the most useful translational models ^3,4^, but at the same time can be more demanding to work with and the tools, time-frame, and costs for genetic manipulation are prohibitive for most studies^5^. Smaller mammals are easier to work with and genetically manipulate, but pose additional challenges in extrapolating findings to humans. One of the most widely prescribed drug classes for prevention of cardiovascular disease in humans, statins, has vastly different effect sizes in rodents as compared to humans^6^. The more rapidly beating hearts of rodents preferentially express fast myofilament and stiff titin isoforms, have increased troponin phosphorylation, and have altered action potential and calcium handling dynamics compared to human hearts^7^. Such protein-level differences can explain why contradictory findings across cardiac studies are common^4^, and why clinical trials of pharmacological interventions can fail to translate effects from animal studies to humans.

Identifying proteins and signaling pathways that are conserved or diverge across commonly used model organisms will contribute to our ability to choose appropriate models for investigating cardiac disease mechanisms and therapies. In addition to varying across species, cardiac protein expression profiles differ across cardiac regions and change in response to disease progression^8^. Mass spectrometry-based methods can assess the species- and regional-specific protein composition of cardiac tissues and can enhance our understanding of the molecular mechanisms underlying the progression of heart disease through proteomic analysis of healthy and diseased samples. This analysis provides a holistic view of the dynamic changes within cardiac proteomes during disease, thereby identifying protein targets relevant to disease^9^.

To facilitate optimal choice of model organism for specific cardiac studies, we set out to perform a systematic comparison of cardiac protein expression in humans and model organisms across the four chambers of the heart. We first utilized a mass spectrometry-based proteomics approach to map the landscape of proteins across the heart and to assess protein abundances in several commonly used model organisms. We then identified protein signatures that are shared or differ across species. We analyzed deep proteomes for the separate cardiac chambers in humans and five relevant model organisms: pig (*Sus scrofa*), horse (*Equus caballus*), rat (*Rattus norvegicus*), mouse (*Mus musculus*) and zebrafish (*Danio rerio*). We identified and quantified ~7,000 proteins in each species and performed quantitative comparisons of protein expression across species with respect to cardiac function and mechanisms of disease. Of those proteins with differential expression between ventricle and atria in all species, a quarter had A-V differential expression that was inverted in some species, reflecting functional differences in heart rate, metabolism, and contractility. Using the differential protein profiles, we show why structural studies of hypertrophic cardiomyopathy are difficult to perform in zebrafish, and we conclude that the best animal model for arrhythmogenic right ventricular cardiomyopathy is pig. These results illustrate how our proteomics resource provides important insights for choice of model organisms in studying disease pathogenesis, ultimately contributing to translatability of findings. This substantial resource quantifying protein abundance in the heart will aid in the discovery of novel molecules and pathways important for cardiac health and disease. Our multispecies cardiac proteome resource is available to the research community as an open-access database: http://atlas.cardiacproteomics.com/. We envision that the content of this database will facilitate experimental design and interpretation of results across species and increase the translational prospect of cardiac findings.

## Results

### Deep proteome profiling of cardiac chambers across six species

To define cardiac protein expression profiles, biopsies from each cardiac chamber from three individuals per species were analysed (Fig. 1A, Supplementary Fig. S1). Specifically, biopsies from left atrium (LA), right atrium (RA), left ventricle (LV) and right ventricle (RV) were collected from three mammals in each group of *Equus caballus* (horse), *Sus scrofa* (pig), *Rattus norvegicus* (rat) and *Mus musculus* (mouse). For *Danio rerio* (zebrafish), atrium (A) and ventricle (V) were collected and pooled from ten fish per sample to ensure sufficient tissue material. For *Homo sapiens* (humans) LA, RA and LV biopsies were taken during mitral valve replacement surgery, where collection procedure via right atrium precluded the possibility to sample from right ventricles. All biopsies were snap frozen in liquid nitrogen immediately after collection and stored at −80°C until further processing. Biopsies were homogenized using a ceramic bead mill and proteins were extracted with a detergent-based buffer, which solubilizes cellular membranes and compartments ^10,11^. Protein extracts were digested into peptides and pre-fractionated at high pH by reverse phase liquid chromatography (RP-HPLC) before mass spectrometry (LC-MS/MS) analysis on a high-resolution Q-Exactive HF quadrupole Orbitrap tandem mass spectrometer ^10,12^. In total, the study covers 654 LC-MS runs amounting to over 40 days of MS measurement time. All 654 raw data files are made available via the Pride repository (see *Data Availability*). In Andromeda data search, only canonical protein sequences were included as global isoform-specific quantification cannot be done accurately by label-free approaches. Despite restraining our search to canonical protein sequences, we measured ~7,000 proteins in each cardiac chamber for each species (Fig. 1B). Evaluation of acquired data is presented in Supplementary Figures S2-S7. For human heart, we measured 6,729 proteins, which are specified in Supplementary Table S1. For mouse heart, we measured 6,943 proteins (Supplementary Table S2), for rat heart we measured 7,446 proteins (Supplementary Table S3), for pig heart we measured 7,177 proteins (Supplementary TableS4), for horse heart we measured 6,479 proteins (Supplementary Table S5), and for zebrafish heart we measured 7,177 proteins (Supplementary Table S6). For each species, we found high correlation between the three biological replicates with Pearson correlation coefficients mostly above 0.9, and principal component analyses showed that in general most of the variance between samples stemmed from differences between cardiac chambers (Supplementary Figures S2-S7). The quantitative proteomics dataset acquired represents a comprehensive mapping of cardiac protein expression profiles across chambers for human heart and five model organisms.

**Figure 1.**
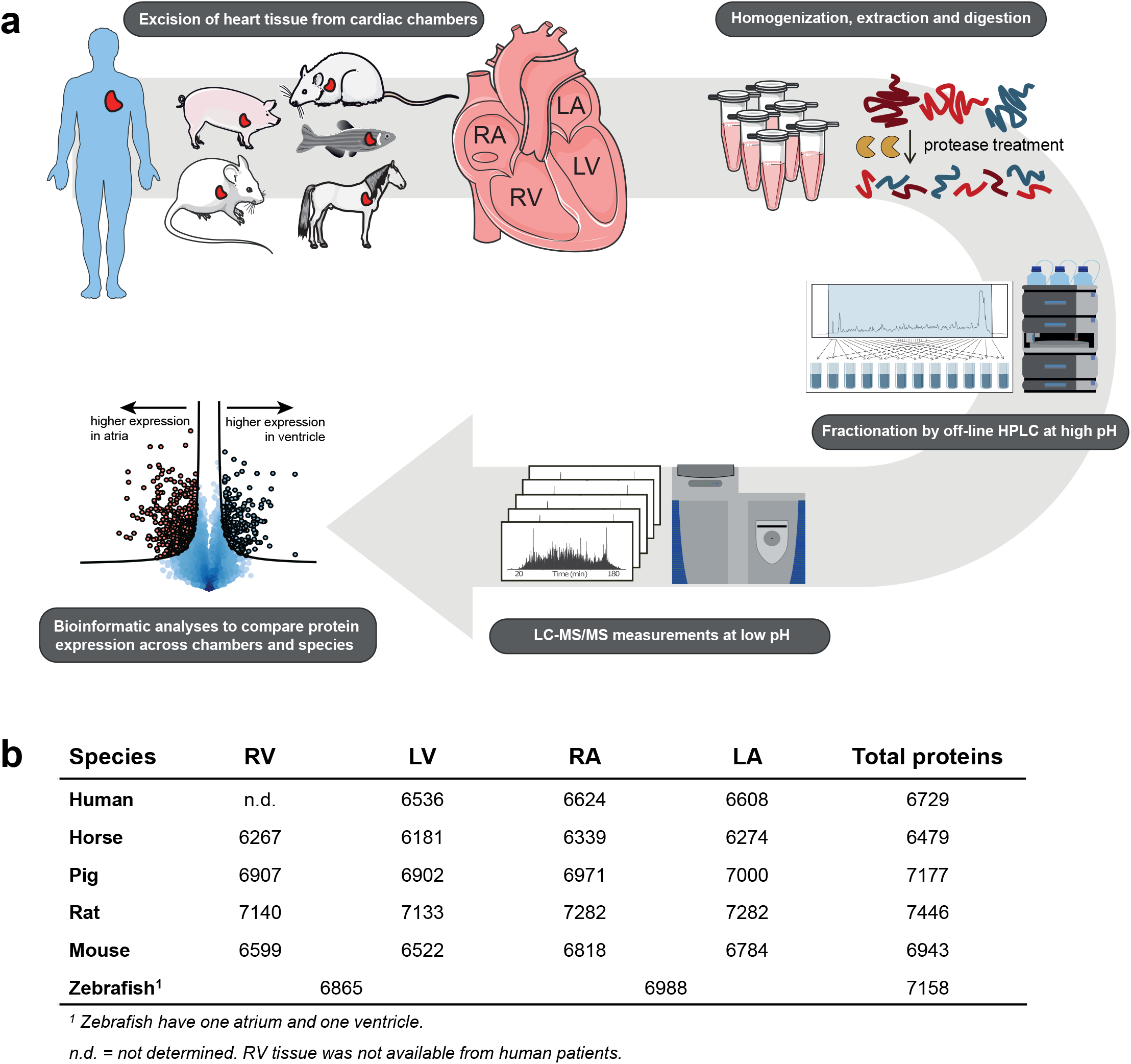
Multispecies proteome mapping across cardiac chambers. a. Workflow for the determination of chamber-specific cardiac proteomes in human, horse, pig, rat, mouse, and zebrafish. Tissue biopsies were collected in triplicates. Biopsies were homogenized followed by protein extraction and digestion, desalted peptides were then fractionated and the generated peptide fractions were analyzed by LC-MS/MS. Data was analysed using MaxQuant and Perseus software. b. Table summarizing the number of proteins measured in each species across cardiac chambers.

### Building a database of protein profiles for all cardiac proteins across species

Differences in cardiac protein profiles across species have direct implications for experimental designs in biomedical studies that aim to identify molecular mechanisms of cardiac disease states; for hypothesis generation as well as for biomarker discovery. To make the large resource of cardiac protein expression data accessible to the broader research community, we aimed to present our knowledgebase of cardiac protein signatures as a database. Since protein homology often creates one-to-many or many-to-one relationships of protein evolution across species, we created a database format that allows to contain these relationships fully and non-redundantly. To this end, we performed protein orthology/paralogy mapping based on EggNOG fine-grained orthology groups ^13,14^, which allowed us to preserve the full information available from our dataset in the database. This protein orthology network contained a total of 34,241 proteins connected by 294,850 binary relationships. To make protein expression differences comparable across species, all raw data was first normalized to a common scale (Supplementary Figure S8, Supplementary Table S7), and protein abundance representations were translated from MS based intensities into a confidence score (Sup. Figure S9)^15^.

We built an open-data knowledgebase of cardiac protein profiles providing easy and efficient access to high quality data in an easy-to-use web interface with intuitive data illustration capabilities with complete overview of protein abundance of orthologues and paralogues across species: http://atlas.cardiacproteomics.com/. The design of the web interface is illustrated in Figure 2. The webpage allows straight-forward comparison of any identified protein in the dataset to all its homologues across species. Any protein can be queried in the online database, and the output returns information on protein expression levels of all orthologues and paralogues across chambers for the analysed species. Specific proteins of interest can thus easily be queried and protein abundance across species and chambers evaluated. This allows individuals to make informed decisions on target proteins, model organisms, and experimental design.

**Figure 2.**
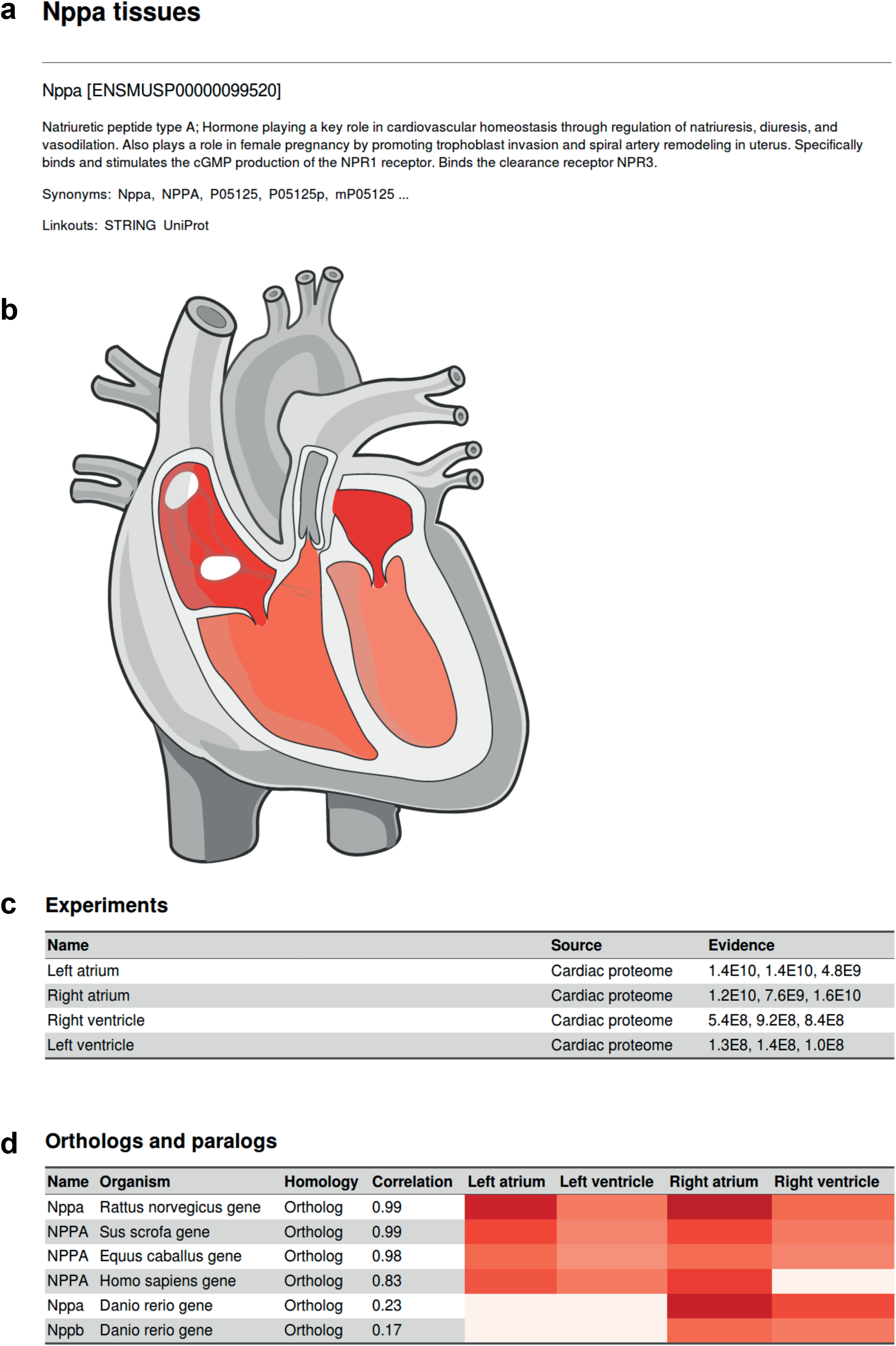
Website interface of cardiac protein expression database across species. Example interface when searching for a protein of interest, example here is natriuretic peptide type A, Nppa, in mouse. **a.** Detailed information of the queried protein as extracted from Uniprot. A link for protein-protein interaction network of the protein as reported in STRING is provided. **b.** Our measured protein expression across mouse heart chambers are displayed on a color scale in a graphic representation of the heart. **c.** Table summarizing the measured experimental data. In this case MS-based intensities were measured in all triplicates from all chambers. The measured protein intensity is provided in the ‘Evidence’ coloumn. Protein abundance is two orders of magnitude greater in the atria than in the ventricles. **d.** All orthologs and paralogs identified in the dataset for Nppa are displayed in an adjacent table for comparison. In the database, measured protein intensitites are translated into a multispecies confidence score for improved comparability.

### Evolutionary conserved cardiac protein profiles

From the quantitative proteomics datasets measured, global comparisons of cardiac protein profiles across species can be achieved. Comparing protein expression profiles across species is not trivial since speciation events have led to a multitude of orthologous and paralogous genes that need to be mapped to their closest relatives. Based on categorizing proteins into orthologous groups using EggNOG^13,14^, as explained above, we created a two-dimensional dataset for visualization by retaining one protein for each species per orthologue group based on highest degree of homology. Unsupervised hierarchical clustering on the resulting orthologue groups (Fig. 3A) showed that i) each species forms a cluster and ii) atria and ventricles form separate clusters. Notably, the species branches clustered according to evolutionary distance, with horse and pig, as well as mouse and rat forming common clusters on species level. As the most distant evolutionary species, the zebrafish had the largest vertical distance in the clustering. Cluster separation by cardiac chamber was particularly clear for smaller mammals and zebrafish, where there were even separate clusters for left and right parts of atria and ventricles. Achieving side-specific cluster separation for atria and ventricle for the smaller mammals is likely a result of these animals being inbred strains and hence posing less molecular heterogeneity. The unsupervised hierarchical clustering underscores that our quantitative proteomics data reflects evolutionary relations between species. To identify essential components of all hearts, we analysed proteins that exhibited similar expression profiles across all species as evaluated by ANOVA analysis. The group of proteins that had similar abundance profiles across all species were analysed by gene ontology enrichment analysis. We found a major overrepresentation of cytoplasmic and mitochondrial proteins as well as proteins involved in translation and metabolic processes (Fig. 3b). These findings are in line with previous reports^16^ and underscore essential characteristics of the heart, such as its high energy demand. Among clusters that were significantly different between species, we found highest enrichment of cytoplasmic, vesicular and mitochondrial proteins, proteins involved in binding and localization, RNA, peptide and small molecule metabolic and catabolic pathways, as well as proteins with structural molecule activity (Fig. 3b).

**Figure 3.**
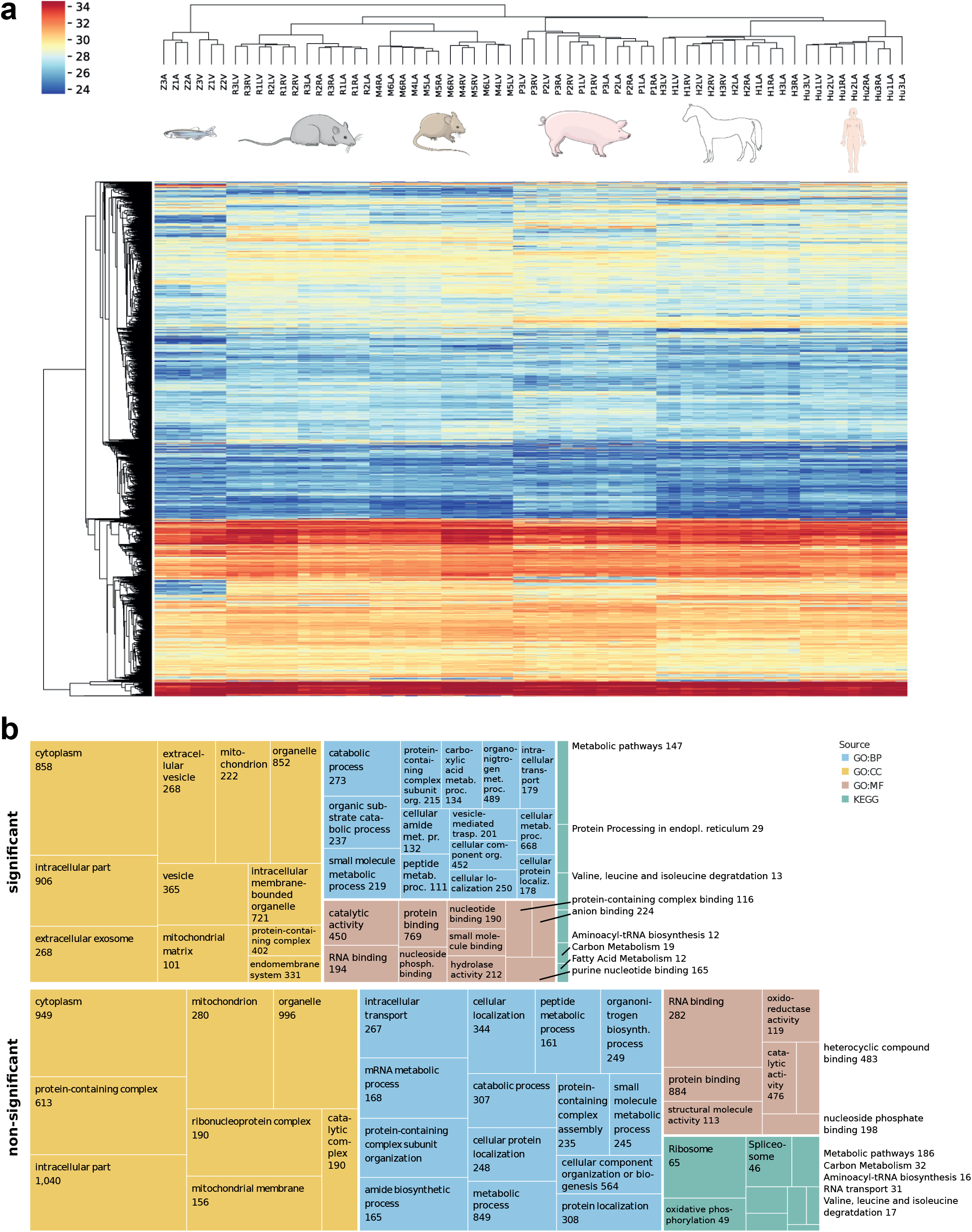
Protein abundance profiles across species. **a.** Unsupervised hierarchical clustering of normalized protein intensities, for proteins measured in all samples resulted in grouping of samples from the same organism and reflects evolutionary distance between species as well as specific similarities and differences in protein expression. Proteins are coloured by intensity with red showing highest and blue lowest intensity values (color bar denotes log2 transformed normalized protein intensities). **b.** Visual representation of gene ontology (GO) enrichment analysis of proteins with significantly different (upper panel) or similar (lower panel) abundance profiles across all species. Shown are representative enriched terms for GO biological process (BP) cellular component (CC), and molecular function (MF), as well as Kyoto Encyclopedia of Genes and Genomes (KEGG) pathways. Sizes of boxes are proportional to –log10(p-value) of the enrichment (the larger the more significant) and numbers denote the number of proteins enriched in the respective category.

### Species-specific differences in heart proteins

We used Principal Component Analysis (PCA) to reduce the dimensions of the protein expression profiles and to define proteins driving differences amongst the cardiac chambers and between species. Most of the variance in the dataset was explained by differences in expression between zebrafish, large mammals and small mammals, which formed separate groups along principal component 1 and 2 (Fig. 4a, upper panel). Prominent proteins driving this differentiation included NPPA, MYL7, MYL4, MYH11, SLC8A1 and ATP2B1, as well as several other channels, myofilaments and extracellular matrix proteins (Fig. 4a, lower panel). Thus, essential molecular elements of fundamental cardiac functions are among the most differentially expressed proteins across species.

**Figure 4.**
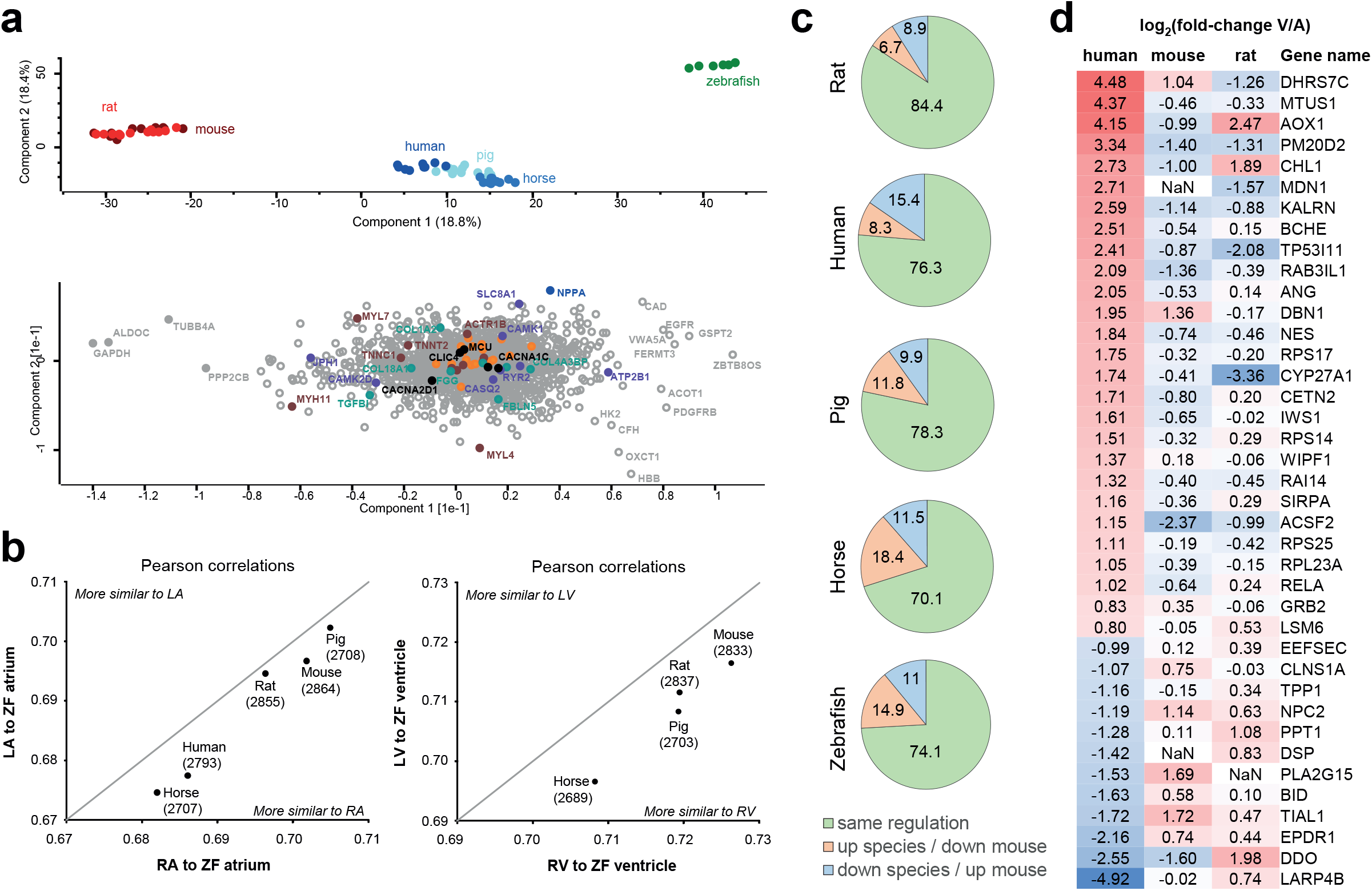
Species-specific differences in protein abundance profiles. **a.** Principal component analysis (PCA, top) shows that the main sources of variation in the dataset is contributed by i) the zebrafish samples (green) being different from all mammalian samples, and ii) small mammals (red) being different from large mammals (blue). Analysis of which proteins explain most of the sample variance between samples (bottom) highlights e. g. MYL4 and MYL7 showing high variance between zebrafish and large mammals along component 2, as well as NPPA, MYH11 and SLC8A1 highlighted as major contributors to the inter-mammalian differences. A reference set of mitochondrial proteins are highlighted in orange, which show comparably lower loadings. **b.** Analysis of zebrafish protein abundance profile in atria and ventricle compared to corresponding mammalian protein abundance profiles in left and right ventricle and atria. Pearson correlation analysis across all proteins consistently shows higher similarity to right side of the mammalian hearts, for ventricle as well as atria. **c.** Comparison of protein abundance differences between atria and ventricles across species. Protein abundances in ventricle compared to atria was calculated for mouse heart and ratios were compared to all other species. Pie charts illustrate the percentage of proteins showing similar protein ratios (green), higher abundance in atria in other species in contrast to mouse (orange) and higher abundance in ventricle in other species contrary to finding in mouse (blue). **d.** Proteins significantly differentially expressed between human ventricle and atria, which show opposite abundance profile in mouse and/or rat compared to human. log2 fold-change of atria versus ventricle are shown, i. e. proteins higher expressed in ventricle are denoted positive (red), those higher expressed in atria are denoted negative (blue).

To explore which cardiac protein functional groups are differentially regulated between species, we performed ontology enrichment analyses on protein expression profile clusters (hierarchical clustering) which were significantly different between species by ANOVA analysis (Sup. Fig. S10). Proteins that have higher expression in small mammals (mouse and rat) compared to large mammals and zebrafish were enriched for mitochondrial proteins as well as proteins involved in ligation, translation and peptide biosynthesis. Proteins that have lower expression in small mammals compared to other species were enriched for cellular amino acid metabolism. Taken together, this may indicate differences in energy metabolism in hearts of small rodents. Several transcription factors were also enriched in both sets of clusters (ZF5, E2F-3, HES-7, Sp1), potentially indicating differential regulation of downstream proteins.

Proteins differentially expressed in human compared to all other species (Sup. Fig. S11) were enriched for sarcolemma, structural constituent of muscle, Sphingolipid pathway, and voltage-gated calcium channel activity. Specifically, ATP1A1, MYH11, JPH1, CACNA1C, CACNB2 and CAMK1 were among the significantly different proteins in human, highlighting that proteins fundamental to cardiac function can be some of the most differentially expressed compared to model organisms. Other proteins with a unique protein profile for the human heart include versican (VCAN), an extracellular proteoglycan involved in heart development; plectin (PLEC), a cytoskeletal linker found in nearly all mammalian cells; and transgelin (TAGLN), an actin binding protein involved in Ca-independent smooth muscle contraction.

Zebrafish is a popular model organism in cardiac research, although its physiology with only two cardiac chambers is markedly different from mammals. We examined which side of the mammalian heart the two-chambered zebrafish heart resembles most with regards to its molecular profile. To this end, we compared protein abundancies between zebrafish and all other species using Pearson correlation, Euclidian distance and Cosine similarity. Our analyses consistently showed greatest similarity between zebrafish heart and the right half of mammalian hearts (Fig. 4B and Supplementary Figure S11). This was the case for atria as well as ventricle. We propose this to reflect the zebrafish circulatory system being a low-pressure system, and hence the function of the heart resembling the right side of mammalian hearts serving the lower-pressure pulmonary system.

Lastly, we compared the protein differential expression between atria and ventricles across all species. We computed protein expression fold-change between atria and ventricles for each species and determined differential expression between the chambers by two-sample t-test. We compared these significantly different proteins from each species to the respective fold-change expression in mouse, as mouse data showed the highest degree of completeness in the EggNOG mapping. This analysis revealed that 70-85% of all proteins showed the same, and 20-25% showed opposite regulation across species (Figure 4C, Figure S10A+B). These proteins with opposing expression patterns include proteins implicated in cardiac function and disease: For instance, we identified 39 proteins that were significantly overexpressed in human atria or ventricle but showed opposite expression patterns in mouse and/or rat (Figure 4D). These proteins included important desmosomal proteins such as desmoplakin (DSP), transcription factors such as NF-kappa-B (RELA) and cytoskeleton-modifying proteins such as MTUS1 (microtubule-associated tumor suppressor 1), drebrin (DBN1) and nestin (NES).

### Molecular assessment of model organisms for cardiac disease studies

In addition to the broader implications of the above analyses on the translation of molecular cardiology studies between species, we wanted to attain a more refined picture of disease-specific differences between species by comparing expression of proteins known to be involved in particular diseases. We compared left ventricular protein expression across all species for proteins involved in hypertrophic- and dilated cardiomyopathy (HCM and DCM) ^17^ and performed unsupervised hierarchical clustering on those proteins using Euclidian distance (Figure 5A+B). As expected, the species clustered according to their evolutionary distance, with human and pig being most closely related, followed by horse, then mouse and rat, and finally zebrafish. Notable differences for the HCM associated genes include lower expression of MYL2 and 3, ACTC1 and MYH7 in zebrafish in comparison to the other species, indicating that extra care has to be taken when translating study results from zebrafish to human for these particular proteins. In DCM, expression of cytoskeletal and contractile proteins such as tropomyosin 1 (TPM1), nebulette (NEBL), troponin 1 (TNNI3), laminin (LAMA2), dystrophin (DMD) and actin (ACTC1) was again lower in zebrafish in comparison to the other species. Considering the crucial functions of these proteins in cardiac muscle tissue, attention has to be paid when designing studies in zebrafish involving these proteins.

**Figure 5.**
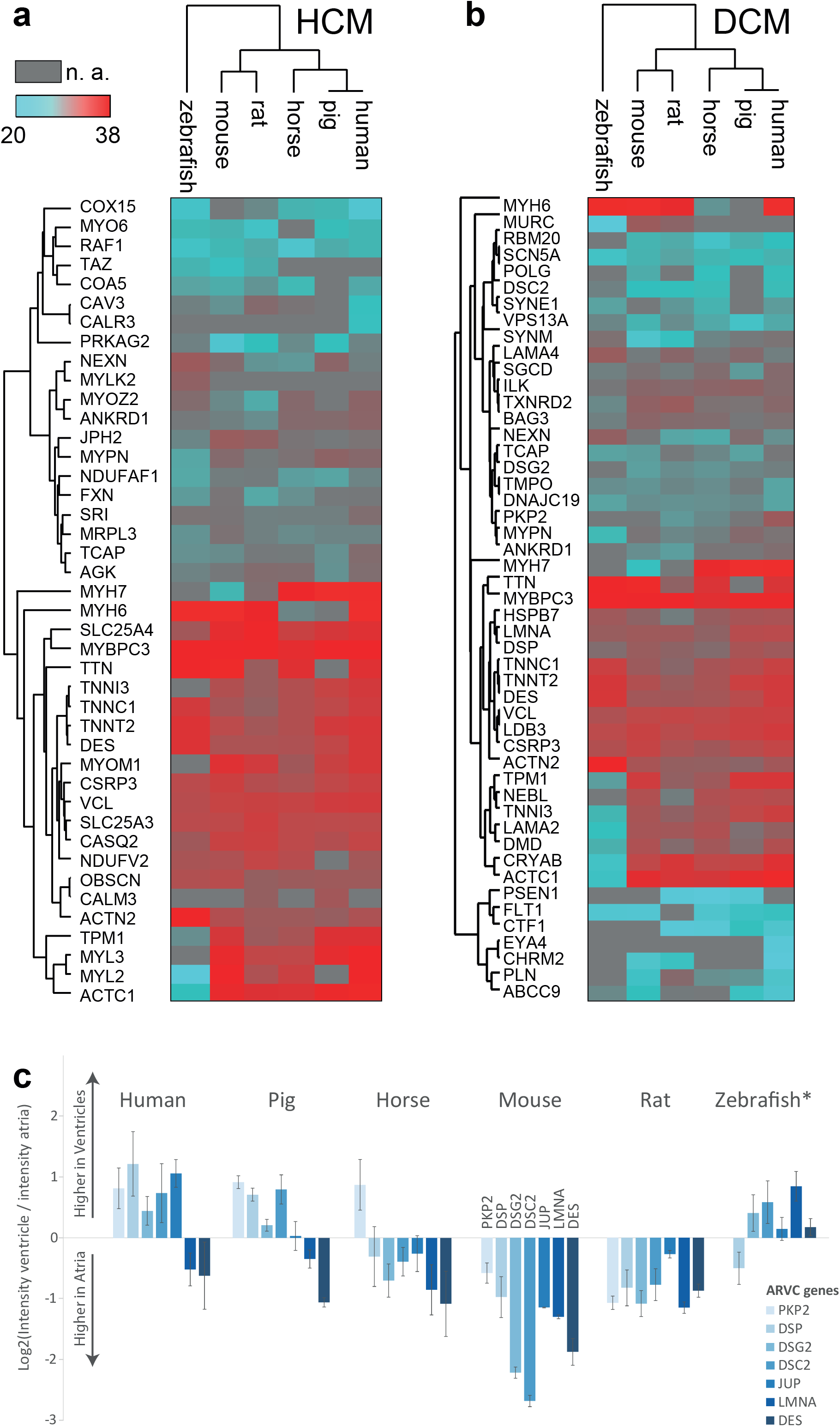
Protein abundance profiles for cardiac disease associated proteins across species. **a+b.** Median protein abundances in left ventricles are shown for protein reported to be involved in hypertrophic cardiomyopathy (HCM, panel a) and dilated cardiomyopathy (DCM, panel b) across species. Color scale represents log2-transformed protein intensities (red – highest abundance, turquoise – lowest abundance, grey – not available). **c.** Comparison of protein abundance ratios between atria and ventricle for proteins encoded by seven genes involved in arrythmogenic right ventricular cardiomyopathy (ARVC). log2 fold-change between ventricle and atria are shown. Note that the human ratio profile is best reflected by pig, while profiles in other species differ markedly.

Finally, we analyzed the expression pattern of the seven proteins most commonly involved in arrhythmogenic right ventricular cardiomyopathy (ARVC) across species ^18^. From a molecular standpoint, the most suitable model organism to study ARVC should show protein expression patterns as similar as possible to human. We found that five of the primary ARVC-associated genes were more highly expressed in the ventricles of human, whereas expression patterns varied considerably among the other species, with the pig ARVC-associated protein expression profile being the most concordant with human (Fig. 5C).

## Discussion

In the current study, we utilized a proteomics approach to generate a high-resolution map of the cardiac protein landscape allowing for comparison of protein abundances across commonly used model organisms. Using this resource, we have shown which protein profiles are shared and which differ across species. Specifically, we present a quantitative high-resolution map of cardiac protein expression across humans and five common model organisms. Previous studies of model organisms have illuminated portions of their cardiac proteomes ^10,19,20^, but often with a particular focus such as cardiac development ^21^, disease models ^22^, subcellular protein expression ^23,24^, phosphorylation ^25–27^, protein turnover ^8,28^ or focused on smaller mammals and amphibians ^16^ or human tissue collected several days post-mortem ^29^. No study has been presented that allowed for direct comparison of cardiac protein profiles between model organisms and human. Gaining insight into the molecular build-up of the human heart in homeostasis and in disease states and directly comparing this build-up to model organisms is essential for selecting appropriate models for investigation of cardiac disease. For most model organisms used in cardiac research, essential information about the molecular composition of the heart is still lacking. Our study closes this knowledge gap with chamber-specific, quantitative information of ~7000 proteins expressed in the hearts of humans and five common model organisms.

Differences in cardiac protein profiles across species have direct implications for experimental design and interpretation of studies that aim to identify molecular mechanisms of cardiac disease, discover relevant biomarkers, or develop pharmacological therapies. The analysis of proteins with differential atrial-ventricular expression in all species studied found that a quarter of these proteins have expression differentials that are inverted in some species compared to mouse. These molecular differences reflect functional differences in heart rate, metabolism, and contractility, and are critical to evaluate prior to choice of model organism. Among proteins that show differential expression between species are proteins vital to cardiac function and causal to diseases such as dilated cardiomyopathy, hypertrophic cardiomyopathy and arrhythmogenic right ventricular cardiomyopathy. While we confirm that the human expression profiles of proteins involved in ARVC are recapitulated in pig hearts, we find that none of the four other model organisms reflect the ARVC protein abundance profile of human hearts. Hence, translating molecular disease mechanisms from these organisms to humans present immediate challenges. This exemplifies the extent to which cardiac protein expression can differ between species for just one major cardiac disease. Yet, we did not find clear or uniform patterns of disease-related differences in protein expression between species beyond protein expression differences being greater with increased evolutionary distance. As demonstrated for DCM and HCM, some proteins showed similar expression across species while other proteins showed opposing profiles. As protein expression differences between species are complex and not predictable for a disease as a whole, the expression of each protein must be individually evaluated, which can be provided by tissue proteomics. The differences in protein abundance outlined in this study, and made accessible via an online database, aid in rationalizing expected translational potential across species.

Comparison across organisms is challenging due to evolutionary distance, but also due to technical aspects of data shortage and knowledge gaps. An important consideration when comparing proteomes across species is the completeness and correctness of employed protein sequence and orthology databases, especially for the less well-studied organisms. These differences could potentially introduce a systematic bias to protein identification and data analysis. Here, we minimize the influence of these systematic differences by employing the gene-centric protein database (ENSEMBL) and the corresponding protein orthology tree (EggNOG). This approach was chosen because the genomes of all investigated species have been sequenced, while knowledge on protein level is vastly different. Here, the employed data-driven approaches such as hierarchical clustering and similarity metrics can yield new insights even when curated knowledge is sparse. As one example, the zebrafish is becoming an increasingly popular model organism in cardiac studies due to its versatile use in high-throughput drug screening, CRISPR technology ^30^ and the possibility to perform *in vivo* optical mapping of action potentials and calcium fluxes ^31^. Querying our quantitative proteome data, we could show that the zebrafish heart is generally more similar to the right side of the mammalian heart, possibly due to their common feature of being a low-pressure system.

We propose that the ability of an animal model to recapitulate human heart disease states is directly related to the similarity in relative abundance of proteins in networks relevant to disease. Here we have integrated experimental data on cardiac protein profiles from humans and five major model systems to generate a resource that facilitates choice of model organism in cardiac disease studies. Our analysis of protein abundance across species revealed molecular features that are shared among all species, as well as specific features that are species dependent, together assembling a portrait of cardiac protein signatures for all commonly used model organisms. This study is the first in-depth, quantitative dataset of cardiac protein expression across humans and common model organisms at cardiac chamber resolution. Our results allow meaningful comparisons both between species as well as between cardiac chambers within a species, even when curated knowledge is sparse. We expect that this data, publicly accessible in database format, will allow cardiac researchers to make informed decisions on experimental design based on the available protein expression data between species. Our data may aid in choosing the best suited model organism to test a given hypothesis, as well as to evaluate findings from studies conducted in model organisms for human physiology.

## Supporting information

Proteins identified in human heart

Proteins identified in mouse heart

Proteins identified in rat heart

Proteins identified in pig heart

Proteins identified in horse heart

Proteins identified in zebrafish heart

Protein intensities for database creation

## Funding

Supported by grants from the Carlsberg Foundation (CF17-0209), The Danish Council for independent Research (DFF-4092-00045), and The Novo Nordisk Foundation (NNF15OC0017586) to AL. The Novo Nordisk Foundation Center for Protein Research is funded in part by a generous donation from the Novo Nordisk Foundation (NNF14CC0001).

## Acknowledgements

Some figure content was downloaded from Servier Medical Art by Servier licensed under a Creative Commons Attribution 3.0 Unported License. The authors would like to thank the PRO-MS Danish National Mass Spectrometry Platform for Functional Proteomics and the CPR Mass Spectrometry Platform for instrument support and assistance.

## Conflict of Interest

None declared.

## >Author contributions

Sample preparation and mass spectrometry measurements (NL, PCP), bioinformatics data analyses (NL, UL, JYZ), construction of database and gold standard evaluation (AS, CS, LJJ), collecting samples and physiological discussions (KC, BHB, MBT, PRL, RWM, MSO). LC-MS/MS supervision (JVO), conception of project and interpretation of data (AL), writing of manuscript (NL, RWM, AL) with approval of content by all authors.

## Materials and Methods

Detailed methods section is provided in the Supplementary Material.

### Materials

If not specified otherwise, chemicals and reagents were acquired from Sigma Aldrich, USA. Chromatography solvents were acquired from VWR, USA.

### Tissue collection

We collected biopsies from left atrium (LA), right atrium (RA), left ventricle (LV) and right ventricle (RV) in mammals and from atrium (A) and ventricle (V) in zebrafish. Human biopsies were collected during minimal invasive mitral valve replacement surgery via the right atrium and due to the nature of this procedure the right ventricle was not accessible and therefore not included. Due to differences in heart sizes, biopsies from human, pig and horse were specifically taken from the muscular part of the free walls, for rat and mouse entire free wall biopsies were collected, and for zebrafish entire chambers were collected and pooled from ten fish per sample. All biopsies were snap frozen in liquid nitrogen immediately after collection and stored at −80C until further processing.

### Tissue homogenization, digestion and fractionation

Frozen tissue biopsies were homogenized on a Precellys24 homogenizer (Bertin Technologies, France) with ceramic beads (2.8 and 1.4mm zirconium oxide beads, Precellys) in tissue incubation buffer (50mM Tris-HCl, pH 8.5, 5mM EDTA, 150mM NaCl, 10mM KCl, 1% Triton X-100, 5mM sodium fluoride (NaF), 5mM beta-glycerophosphate, 1mM Na-orthovanadate, containing Roche complete protease inhibitor). After homogenization, samples were incubated for 2h at 4°C (20rpm). Samples were centrifuged (15000x g, 20min, 4°C) and the soluble fraction was collected and protein precipitated using ice-cold acetone (25% final concentration, VWR, USA) for 1h at - 20°C followed by centrifugation (400x g, 1.5min). Supernatants were discarded and protein resuspended in Guanidine-HCl buffer (6M Gnd-HCl, 50mM Tris-HCl, pH 8.5, 5mM NaF, 5mM beta-glycerophosphate, 1mM Na-orthovanadate, containing Roche complete protease inhibitor, 5mM Tris(2-carboxyethyl)phosphine (TCEP), 10mM chloroacetamide (CAA)) and incubated in the dark at room temperature (RT) for 15min. Protein was digested using endoproteinase Lys-C (Trichem ApS, Denmark; 1:100 w/w) for 1h, 750rpm at 30°C in the dark, followed by dilution (1:12 with 50mM Tris-HCl, pH8) and digestion with trypsin overnight (16h) at 750rpm and 37°C (Life technologies, USA, 1:100 w/w). Digestions were quenched by addition of trifluoroacetic acid (TFA, 1% final conc.) and centrifuged (14000x g, 10min). Soluble fractions were desalted and concentrated on C18 SepPak columns (Waters, USA) according to manufacturer’s protocol. Up to 1mg peptide was fractionated by reverse-phase high pressure liquid chromatography (HPLC) on an Dionex UltiMate 3000 HPLC system (Thermo Scientific, USA) equipped with an XBridge® BEH C18 Sentry Guard Cartridge pre-column (130Å, 3.5um particle size, 4.6*20mm, Waters, USA) coupled to an XBridge® Peptide BEH C18 packed column (130Å, 3.5um particle size, 4.6*250mm, Waters, USA) at 1mL/min flow rate. The following gradient elution program was used at a constant supply of 10% solvent C (25mM ammonia, pH10): 0-49 min: 10-25% solvent B (100% ACN) linear gradient, 50-54 min: 25-70% B linear gradient, 55-59 min: 70% B isocratic flow, followed by column re-equilibration at 5% B for 10min as previously described ^12^. Peptides were collected from 0-60 minutes in 10 concatenated fractions. Fraction volume was reduced by vacuum centrifugation to 20-100µL.

### LC-MS/MS measurements

Fractionated peptide samples were analyzed by online reversed-phase liquid chromatography coupled to an Q-Exactive Plus quadrupole Orbitrap tandem mass spectrometer (Thermo, Bremen, Germany). Peptide samples were separated on 15cm fused-silica emitter columns pulled and packed in-house with reversed-phase ReproSil-Pur C18-AQ 1.9um resin (Dr. Maisch GmbH, Ammerbuch-Entringen, Germany) in a 1h multi-step linear gradient (0.1% FA constant; 2-25% ACN in 45min, 25-45% ACN in 8min, 45-80% ACN in 3min) followed by a short column re-equilibration (80-5% ACN in 5min, 5% ACN for 2min).

Raw MS data was processed using the MaxQuant software (version 1.5.3.19, Max-Planck Institute of Biochemistry, Department of Proteomics and Signal Transduction, Munich, Germany) and proteins identified with the built-in Andromeda search engine based on ENSEMBL^32^ canonical protein collections for each species. False-discovery rate cutoffs were set to 1% on peptide, protein and site decoy level (default), only allowing high-quality identifications to pass. Because all raw intensities showed similar distributions, data was normalized across species by quantile normalization based on the Bioconductor R package LIMMA^33^.

We normalized the data globally across species by median centering and used the EggNOG database^13^ to map orthologous groups of proteins between species. We then systematically compared similarities and differences in protein expression across species. Data analysis was be performed using Perseus^34^, Cytoscape^35^, R and Python. For representation in the database, the intensity values were translated into a multispecies confidence score by comparison to a gold standard as previously described^15^. See further details in the supplementary methods section.

### Data availability

The mass spectrometry proteomics data have been deposited to the ProteomeXchange Consortium via the PRIDE^36^ partner repository with the dataset identifier PXD012636 (accessible through https://www.ebi.ac.uk/pride/archive/login, username, password: CW6iz45X) and project name ‘The protein expression landscape of the heart across humans and model organisms’. The website containing all cardiac protein expression data across species is accessible under atlas.cardiacproteomics.com

## Notes

https://atlas.cardiacproteomics.com/

