## Supplementary Material for "Proteomics Dissection of Cardiac Protein Profiles of Humans and Model Organisms"

### Supporting Information

### Materials and Methods

#### *Tissue collection*

All animal experiments were performed according to the European Union legislation for protection of animals used for scientific experiments and was approved by the Danish National Animal Experiments Inspectorate. Housing facilities followed the Federation of European Laboratory Animal Science Associations (FELASA) guidelines for health monitoring, testing and veterinary care (www.felasa.eu).

*Zebrafish*

Adult wildtype AB genotype zebrafish (all siblings, bred in own facility) were kept at 28.5$^{o}$C and standard housing facilities. The zebrafish were euthanized by rapid freezing in ice water and the hearts were quickly excised and washed in Ringer´s solution (116mM NaCl, 2.9mM KCl, 1.8mM $CaCl_2$, 5mM HEPES, pH 7.2). The two cardiac chambers were dissected, snap-frozen in liquid nitrogen and stored at -80$^{o}$C until use. Atria and ventricles from 10 zebrafish were pooled for each biological replicate. In total, 30 zebrafish were dissected to yield three atria and three ventricle samples. Experiments with the zebrafish were performed under license 2012-14-2934-00041.

*Mouse*

Three male C57BL6 BomTac mice at an age of 8 weeks were purchased from Taconic (Ry, Denmark) and acclimatized for one week before starting experiments. The mice were housed at a constant room temperature of 22$^{o}$C and had access to water and food *ad libitum*. The mice were euthanized by cervical dislocation and the hearts were explanted, washed in 0.9% NaCl and dissected into the four chambers. The cardiac chambers were immediately snap-frozen in liquid nitrogen and stored at -80$^{o}$C until use.

*Rat*

Three male Wistar rats (Taconic, Ry, Denmark) with a weight of 150g were acclimatized one week prior to starting the experiment. The rats were housed at a constant room temperature of 22$^{o}$C and had access to water and food *ad libitum*. They were euthanized by cervical dislocation. The thorax on the rats was opened, the heart explanted and quickly washed in 0.9% NaCl. The four chambers were dissected, immediately frozen in liquid nitrogen and stored at -80$^{o}$C until use.

*Pig*

Three 11 weeks old (25-35kg) Danish landrace pigs (sows) were anesthetized by intramuscular infusion of zoletil pig mixture (1mL/10 kg; 250mg dry tiletamin+zolazepam, 6.5mL xylazine 20mg/mL, 1.25 ketamine 100mg/mL, 2.5mL butorphanol 10mg/mL, and 2mL methadone 10mg/mL). The pigs were moved to the operating theater, euthanized with pentobarbital (intravenous), and the hearts were removed via thoracotomy and placed in an ice-cold cardioplegic solution (110mM NaCl, 16mM KCl, 16mM $MgCl_2$, 1.2mM $CaCl_2$, 10mM $NaHCO_3$).

The left and right atrial appendages were removed and transmural samples were collected from the middle portion. Middle anterior transmural sections from the left and right ventricle, as well as samples from the mid-myocardium were collected. Tissue isolation was performed at $4^oC$ and samples were rinsed in cold phosphate-buffered saline (PBS) before snap freezing in liquid nitrogen. All tissue sections were stored at $-80^oC$ until use.

All pig experiments were performed under a license from the Danish Ministry of Environment and Food (license No. 2012-15-2934-00083) and in accordance with the Danish guidelines for animal experiments according to the European Commission Directive 86/609/EEC license No. 2012-15-2934-00083. Pigs were acquired from Krigsagergård w. Henrik Larsen, Gilleleje, Denmark.

*Horse*

Biopsies from each of the four cardiac chambers were obtained from three horses. The horses were donated to the Department of Veterinary and Animal Sciences, University of Copenhagen (Copenhagen, Denmark) and were all euthanized by captive bolt gun and bleeding. To collect equine tissue no permit for animal testing were necessary under Danish law (The Animal Experimentation Act 1253 of $8^{th}$ of March 2013). All horses were mares, one eight-year-old tinker and two 16-18 years old small mixed breeds. The prevalent reason for euthanasia was lameness, the horses had no history of any cardiac illness. Tissue samples from the left and right atria (pectinate muscles, mid-myocardial region) and from the left and right ventricle (apical free wall, mid-myocardial region) was sampled right after bleeding, rinsed in cardioplegic solution (129mM NaCl, 12mM KCl, 0.9mM $NaH_2PO_4$, 20mM $NaHCO_3$, 1.8mM $CaCl_2$, 0.5mM $MgSO_4$, 5.5mM glucose, pH 7.4, 4˚C) before snap freezing in liquid nitrogen and stored at $-80^oC$ until use.

*Human*

Tissue samples were collected from three patients undergoing mitral valve surgery where the samples were collected by the surgeon during surgery just prior to the valve plasty or removal. The samples were immediately snap-frozen in liquid nitrogen and stored at -80°C until use. The biopsies were collected from the lateral wall of the right atrium, the posterior wall of the left atrium and from a papillary muscle in the left ventricle. The patients were 52-64 years old, all provided a written informed consent, were in sinus rhythm and did not have any congenital heart disease. Collection of the human heart tissue for the study was approved by the Ethics Committee of the Capital Region of Copenhagen (protocol reference number: 16238) and was in accordance with the Declaration of Helsinki.

##### *Tissue homogenization, digestion and fractionation*

Frozen tissue biopsies were homogenized on a Precellys24 homogenizer (Bertin Technologies, France) with ceramic beads (2.8 and 1.4mm zirconium oxide beads, Precellys) in tissue incubation buffer (50mM Tris-HCl, pH 8.5, 5mM EDTA, 150mM NaCl, 10mM KCl, 1% Triton X-100, 5mM sodium fluoride (NaF), 5mM beta-glycerophosphate, 1mM Na-orthovanadate, containing Roche complete protease inhibitor). After homogenization, samples were incubated for 2h at 4°C (20rpm). Samples were centrifuged (15000x g, 20min, 4°C) and the soluble fraction was collected and protein precipitated using ice-cold acetone (25% final concentration, VWR, USA) for 1h at -20°C followed by centrifugation (400x g, 1.5min). Supernatants were discarded and protein resuspended in Guanidine-HCl buffer (6M Gnd-HCl, 50mM Tris-HCl, pH 8.5, 5mM NaF, 5mM beta-glycerophosphate, 1mM Na-orthovanadate, containing Roche complete protease inhibitor, 5mM Tris(2-carboxyethyl)phosphine (TCEP), 10mM chloroacetamide (CAA)) and incubated in the dark at RT for 15min. Protein was digested using endoproteinase Lys-C (Trichem ApS, Denmark; 1:100 w/w) for 1h, 750rpm at 30°C in the dark, followed by dilution (1:12 with 50mM Tris-HCl, pH8) and digestion with trypsin overnight (16h) at 750rpm and 37°C (Life technologies, USA, 1:100 w/w). Digestions were quenched by addition of trifluoroacetic acid (TFA, 1% final conc.) and centrifuged (14000x g, 10min). Soluble fractions were desalted and concentrated on C18 SepPak columns (Waters, USA) according to the manfacturer's instructions. Up to 1mg peptide was fractionated by reverse-phase high pressure liquid chromatography (HPLC) on an Dionex UltiMate 3000 HPLC system (Thermo Scientific, USA) equipped with an XBridge® BEH C18 Sentry Guard Cartridge pre-column (130Å, 3.5um particle size, 4.6*20mm, Waters, USA) coupled to an XBridge® Peptide BEH C18 packed column (130Å, 3.5um particle size, 4.6*250mm, Waters, USA) at 1mL/min flow rate. The following gradient elution program was used at a constant supply of 10% solvent C (25mM ammonia, pH10): 0-49 min: 10-25% solvent B (100% ACN) linear gradient, 50-54 min: 25-70% B linear gradient, 55-59 min: 70% B isocratic flow, followed by column re-equilibration at 5% B for

10min as previously described [1]. Peptides were collected from 0-60 minutes in 10 concatenated fractions. Fraction volume was reduced by vacuum centrifugation to 20-100µL.

##### *LC-MS/MS measurements*

Fractionated peptide samples were analyzed by online reversed-phase liquid chromatography coupled to an Q-Exactive Plus quadrupole Orbitrap tandem mass spectrometer (Thermo, Bremen, Germany). Peptide samples were diluted to a concentration of 0.2µg/µL in 5% ACN, 0.1% TFA in 96-well microtiter plates and autosampled (5µL injection volume) into an Easy-nLC system (Proxeon Biosystems, Odense, Denmark). Peptide samples were separated on 15cm fused-silica emitter columns pulled and packed in-house with reversed-phase ReproSil-Pur C18-AQ 1.9um resin (Dr. Maisch GmbH, Ammerbuch-Entringen, Germany) in a 1h multi-step linear gradient (0.1% FA constant; 2-25% ACN in 45min, 25-45% ACN in 8min, 45-80% ACN in 3min) followed by a short column re-equilibration (80-5% ACN in 5min, 5% ACN for 2min). Peptides were ionized using a nano-electrospray ionization source, the mass spectrometer was operated in positive ionization mode. MS spectra (375-1500 m/z) were acquired after accumulation of 3E6 ions in the Orbitrap (maximum fill time of 25ms) at 120,000 resolution. A data-dependent Top12 method then sequentially isolated the most intense precursor ions (up to 12 per full scan) for higher-energy collisional dissociation (HCD) in the octopole collision cell. MS/MS spectra of fragment ions were recorded at a resolution of 30,000 after accumulation of 1E6 ions in the Orbitrap (maximum fill time of 45ms).

##### *Raw data processing*

Raw MS data was processed using the MaxQuant software (version 1.5.3.19, Max-Planck Institute of Biochemistry, Department of Proteomics and Signal Transduction, Munich, Germany) and proteins identified with the built-in Andromeda search engine. Raw MS/MS data from each species was searched against an *in-silico* tryptic digest of a database containing all canonical ENSEMBL[2] protein entries for this species (EggNOG v.5/ENSEMBL - Release-77, (human on version 38), Nov. 2014). The search was performed with Carbamidomethyl as a fixed modification, as well as oxidation (M), acetylation of protein N-termini, deamidation (NQ) and Gln->pyro-Glu as variable modifications. A maximum of two missed cleavages and six variable modifications was allowed. The minimum peptide length was set to 7 amino acids (default) and minimum Andromeda score required for modified peptides was 25, with minimum delta score of 6 (default). Due to the similarity of the samples within each species the match-between-runs option was enabled with default parameters. False-discovery rate cutoffs were set to 1% on peptide, protein and site decoy level (default), only allowing high-quality identifications to pass. Because all raw intensities showed similar distributions, data was normalized across species by quantile normalization based on the Bioconductor R package LIMMA [3].

##### *Cardiac proteomics database across species*

The raw intensity distributions of the six species showed a similar overall shape with only slightly varying median values (Supplementary Figure S8A-B). It was thus sufficient to normalize raw intensity measurements across species by median centering to attain comparable sample distributions (Supplementary Figure S8C). Other normalization techniques such as z-scoring and quantile normalization were also evaluated, but did not improve the result further. Since the original median for all samples was located around 1E8, this value was set as the new global median value for all distributions to retain an interpretable data scale

We used the EggNOG database[4] to map orthologous groups of proteins between species. For representation in the database, the intensity values were translated into a multispecies confidence score by comparison to a gold standard as previously described[5]. Given that the protein intensity distributions were comparable across species, we used the human dataset to make this comparison and calculate the new scoring scheme across species.

To convert intensities into confidence scores, the agreement with the gold standard was quantified using fold enrichment. To calculate fold enrichment, we sorted proteins by intensity value and within sliding windows (window-size=50) calculated the fraction of proteins in the dataset found annotated to heart in the gold standard (derived from UniProt[6] annotations) divided by the fraction expected when randomly sampling proteins from the gold standard. Then, we used the resulting curve (protein intensity, fold enrichment) to fit a function to translate intensities into confidence scores:

[illegible] ,

where x is the mean intensity within a sliding window of 50 proteins (Sup. Figure S9).

Further data analysis was be performed using Perseus [7], Cytoscape[8], R and python.

##### *Mapping of ENSEMBL to EggNOG IDs*

In order to analyze protein expression across species, we mapped all proteins to ortholog groups through the EggNOG database[9,10]. Each such EggNOG cluster (NOG group) may contain more than one protein from a given species, and proteins may be connected to each other through orthologue or paralogue homology. In order to consolidate a one-to-one homology protein cluster used for the analysis across species, the NOG clusters were reduced to contain one protein per species, choosing the sub-cluster with the highest number of orthologue or paralogue links between its members. In cases where two sub-clusters included equal numbers of orthologue or paralogue connections between proteins, the sub-cluster with the highest summed protein intensity across species was chosen.

##### *Hierarchical clustering*

Unsupervised hierarchical clustering of cardiac protein expression across humans and model organisms was based on protein intensities normalized to a common median (1E8) and performed in Python. Protein intensities were log2 transformed and Euclidian distances calculated. Clustering was performed on 1,838 orthologous groups after filtering for 100% valid values, and minimum 1 unique peptide per protein.

Significantly different protein expression between species was determined in Perseus as follows. Median intensity values were computed for each chamber in each species, after which proteins with less than two valid values in each species and chamber were discarded (2238 proteins remaining). Less than 10 missing values were present per sample, for which values were imputed with standard settings in Perseus from a left-shifted normal distribution (scaling to 0.3 standard deviations and 1.8 down shift of original distribution, individually for each sample). Significantly different expression was determined between evolutionary groups of zebrafish, small mammals (mouse and rat) and large mammals (human, pig and horse) by multiple-sample ANOVA. Multiple hypothesis correction was applied by permutation-based false discovery rate cutoff at 0.01 and $S_0$=0.3 and 500 randomizations. Ontology Enrichment analyses were performed in gProfiler[11] (https://biit.cs.ut.ee/gprofiler/gost) and visualized in Tableau (https://public.tableau.com/en-us/s/).

Proteins significantly differentially expressed in human compared to all other species were determined from protein intensities after filtering for at least 2 valid values in each chamber and species (2096 proteins remaining), followed by imputation as stated above (maximum of 25 missing values per sample), and two-sample T-test combined with a permutation-based false discovery rate of 0.01 and S0=0.5 and 500 randomizations.

##### *Study limitations*

To investigate whether the quality of our data is sufficient to perform meaningful statistical analyses, we investigated the comparability of our data between cardiac chambers as well as species. This was necessary because the tissue biopsies were collected from different regions of the heart, and did not originate from age- and sex-matched individuals for the large mammals. Due to the sheer size of the horse and pig heart, it was not possible to utilize a whole cardiac chamber, but rather a small part was dissected. In humans, it was obviously not possible to obtain large biopsies because the samples were not collected *post mortem* but *in vivo*. Hence, we expected higher statistical power for the analyses in mouse, rat and zebrafish where (a) the animals were age- and sex-matched and (b) the whole cardiac chamber of interest was dissected and used for LC-MSMS analysis.

##### *Data availability*

The mass spectrometry proteomics data have been deposited to the ProteomeXchange Consortium via the PRIDE[12] partner repository with the dataset identifier PXD012636 (accessible through https://www.ebi.ac.uk/pride/archive/login, username:, password: CW6iz45X) and project name 'The protein expression landscape of the heart across humans and model organisms'.

### SUPPLEMENTARY TABLES

**Supplementary Table S1: All Proteins identified in human heart tissue.** Proteomic investigation of human heart biopsies from right atria (RA), left atria (LA) and left ventricle (LV) resulted in identification of 6,729 proteins.

**Supplementary Table S2: All Proteins identified in mouse heart tissue.** Proteomic investigation of mouse heart biopsies from right atria (RA), left atria (LA), right atria (RA) and left ventricle (LV) resulted in identification of 6,943 proteins.

**Supplementary Table S3: All Proteins identified in rat heart tissue.** Proteomic investigation of rat heart biopsies from right atria (RA), left atria (LA), right ventricle (RV) and left ventricle (LV) resulted in identification of 7,446 proteins.

**Supplementary Table S4: All Proteins identified in pig heart tissue.** Proteomic investigation of pig heart biopsies from right atria (RA), left atria (LA), right ventricle (RV) and left ventricle (LV) resulted in identification of 7,177 proteins.

**Supplementary Table S5: All Proteins identified in horse heart tissue.** Proteomic investigation of horse heart biopsies from right atria (RA), left atria (LA), right ventricle (RV) and left ventricle (LV) resulted in identification of 6,479 proteins.

**Supplementary Table S6: All Proteins identified in zebrafish heart tissue.** Proteomic investigation of zebrafish samples from atria (A) and ventricle (V) resulted in identification of 7,158 proteins.

**Supplementary Table S7: Protein intensity input file for database.** Protein intensities measured across all species with median subtraction for normalization. Input intensities used in the online database at atlas.cardiacproteomics.com.

### Supplementary Figures

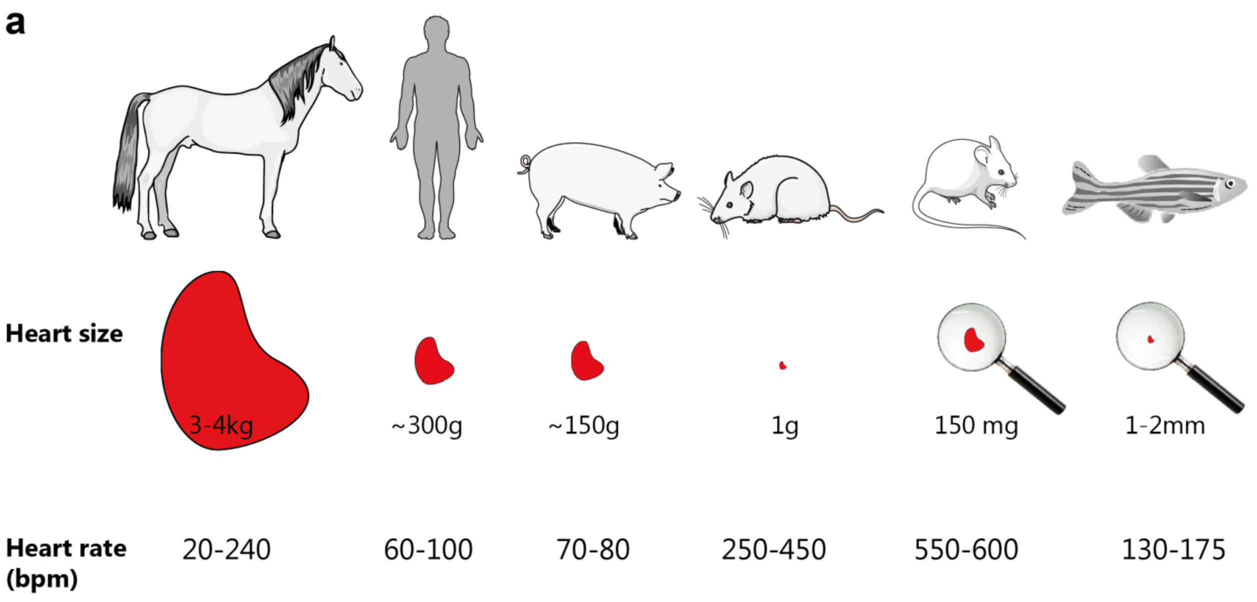

**b**

| No. | Sex | Age at inclusion | Control | NYHA | BMI | Smoking | Alcohol |
|---|---|---|---|---|---|---|---|
| 1 | Male | 52 | yes | 1 | 26.0 | never | 12 |
| 2 | Male | 62 | yes | 1 | 23.0 | never | 3 |
| 3 | Male | 64 | yes | 2 | 26.9 | previously, 22 pack years | 10 |

| No. | Degree of LA dilation | CAD | Hypertension | Diabetes | PVD | Stroke | Medication |
|---|---|---|---|---|---|---|---|
| 1 | severe | no | no | no | no | no | none |
| 2 | severe | no | no | no | no | no | none |
| 3 | severe | no | no | no | no | no | Simvastatin and magnyl |

**Figure S1. Information on samples included in the study a.** Comparison of heart size and heart rate in horse, human, pig, rat, mouse, and zebrafish. The size of the mammal hearts are correlated with their physical size, while the heart rate is negatively correlated. The zebrafish deviates from this trend by having the smallest heart, and only half the heart rate of a rat. Due to differences in heart size, biopsies were collected in different fashions. In humans, biopsies were collected by needle biopsy during cardiac surgery. In horse and pig, biopsies were alike collected from the free walls of the myocardium. In rodents, whole free walls of cardiac chambers were collected. In zebrafish, ten whole chambers were collected and pooled per sample. **b.** Patient information for the three human individuals included in the study. Biopsies were collected from three males undergoing mitral valve replacement surgery. *NYHA*: New York Heart Association functional classification, *BMI*: body mass index, *CAD*: coronary artery disease, *PVD*: peripheral vascular disease, alcohol in units per week.

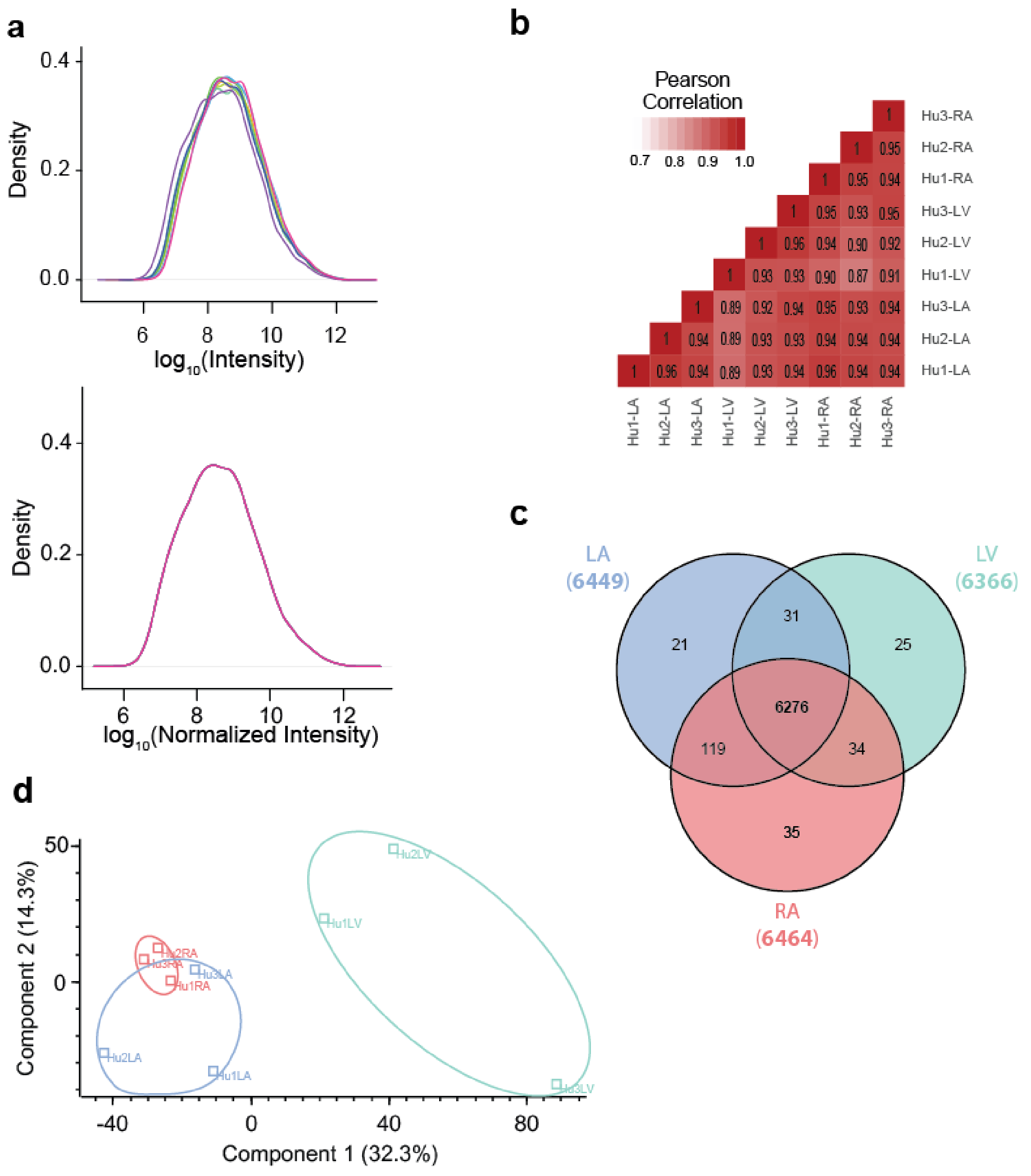

**Figure S2. Evaluation of Human Proteome Data. a.** Density plots of intensities in log10-space displayed before (upper panel) and after quantile normalization (lower panel). Quantile normalization was performed to remove minor technical variation from the dataset. **b.** Pearson correlations coefficients of log-transformed quantile normalized protein intensities across all samples. Hu1 through Hu3 denotes the three human patients in the study, RA, LA, and LV denotes right atrium, left atrium, and left ventricle respectively. **c.** Overlap of identified proteins across heart chambers are shown in a Venn diagram. More than 95% of proteins were identified in all chambers. **d.** Principal component analysis of samples shows clear distinction between atria and ventricle samples in the first principal component that explains 32.3% of the variance in the dataset.

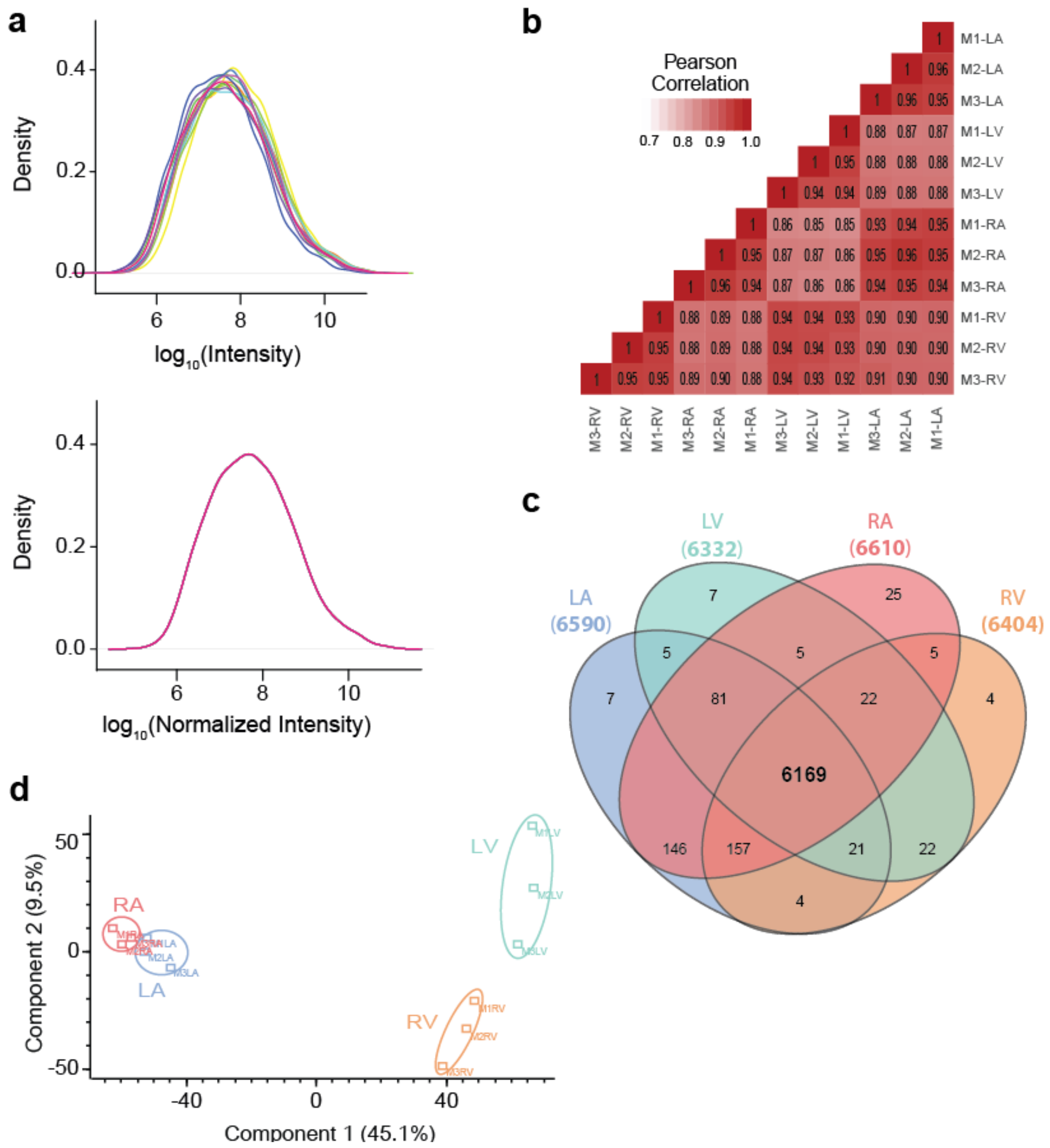

**Figure S3. Evaluation of Mouse Proteome Data. a**. Density plots of intensities in log10-space displayed before (upper panel) and after quantile normalization (lower panel). **b.** Pearson correlations coefficients of log-transformed quantile normalized protein intensities across all samples. M1 through M3 denotes the three mice in the study, LA, RA, LV, and RV denotes left atrium, right atrium, left ventricle, and right ventricle respectively. **c.** Overlap of identified proteins across heart chambers are shown in a Venn diagram. More than 92% of proteins were identified in all chambers. **d.** Principal Component Analysis of samples show clear distinction between atria and ventricle samples, and separation between right and left atria/ventricle in the first two principal components that explain 54.6% of the variance in the dataset.

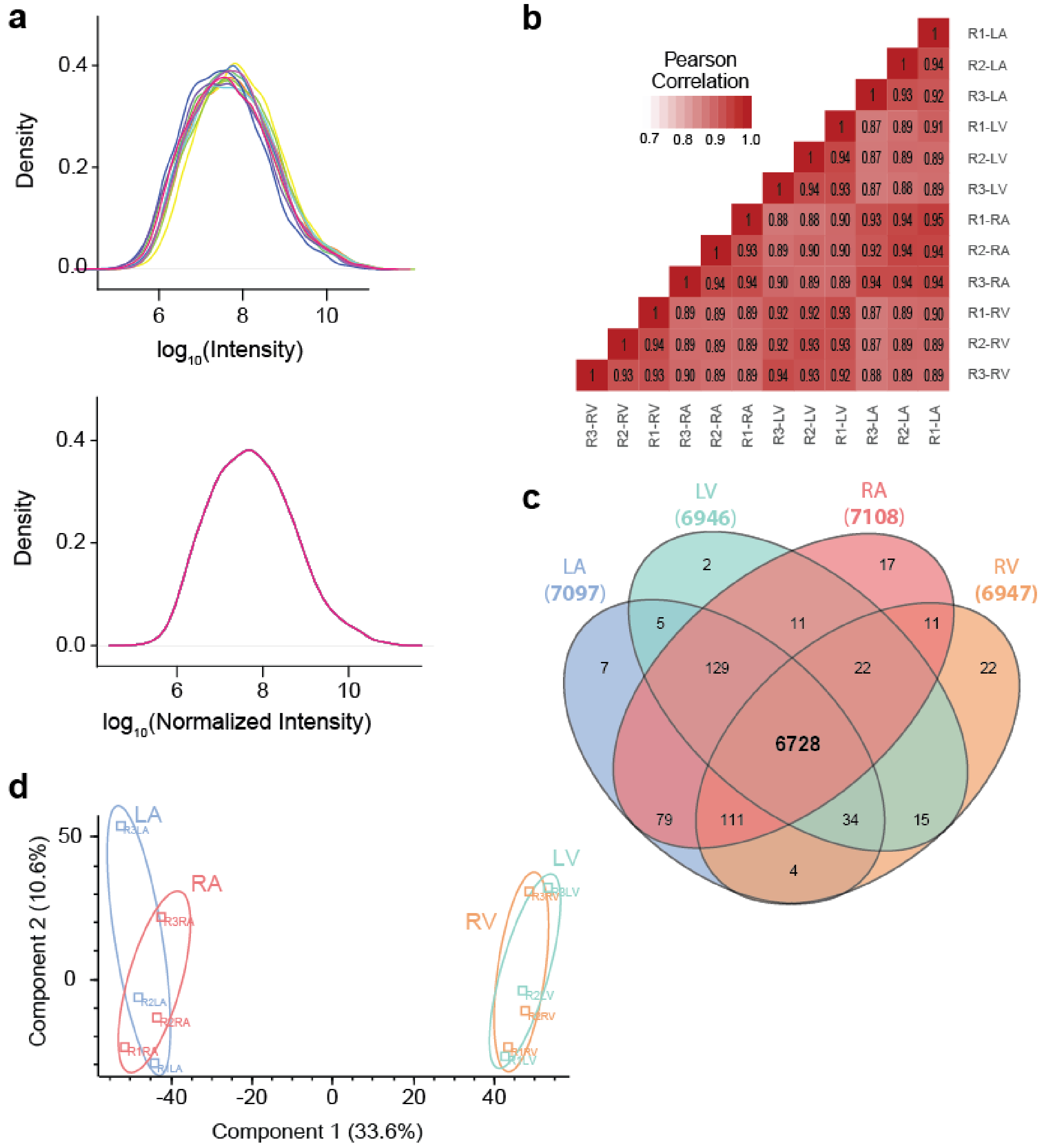

**Figure S4. Evaluation of Rat Proteome Data. a**. Density plots of intensities in log10-space displayed before (upper panel) and after quantile normalization (lower panel). **b.** Pearson correlations coefficients of log-transformed quantile normalized protein intensities across all samples. R1 through R3 denotes the three rats in the study, LA, RA, LV, and RV denotes left atrium, right atrium, left ventricle, and right ventricle respectively. **c.** Overlap of identified proteins across heart chambers are shown in a Venn diagram. More than 93% of proteins were identified in all chambers. **d.** Principal Component Analysis of samples show clear distinction between atria and ventricle samples in the first principal components that explains 33.6% of the variance in the dataset.

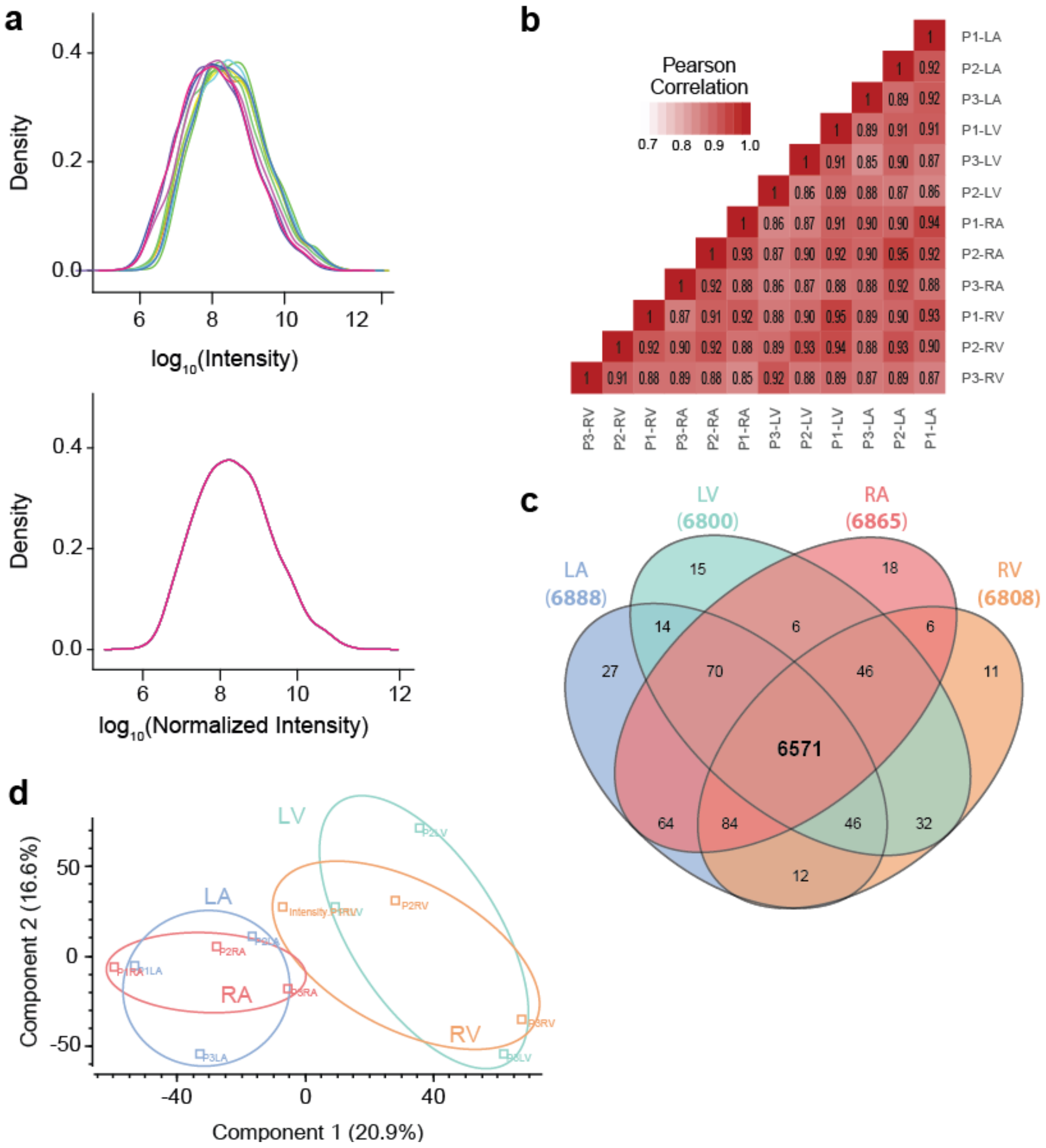

**Figure S5. Evaluation of Pig Proteome Data. a.** Density plots of intensities in log10-space displayed before (upper panel) and after quantile normalization (lower panel). **b.** Pearson correlations coefficients of log-transformed quantile normalized protein intensities across all samples. P1 through P3 denotes the three pigs in the study, LA, RA, LV, and RV denotes left atrium, right atrium, left ventricle, and right ventricle respectively. **c.** Overlap of identified proteins across heart chambers are shown in a Venn diagram. More than 93% of proteins were identified in all chambers. **d.** Principal Component Analysis of samples between atria and ventricle samples in the first two principal components that explain 37.5% of the variance in the dataset.

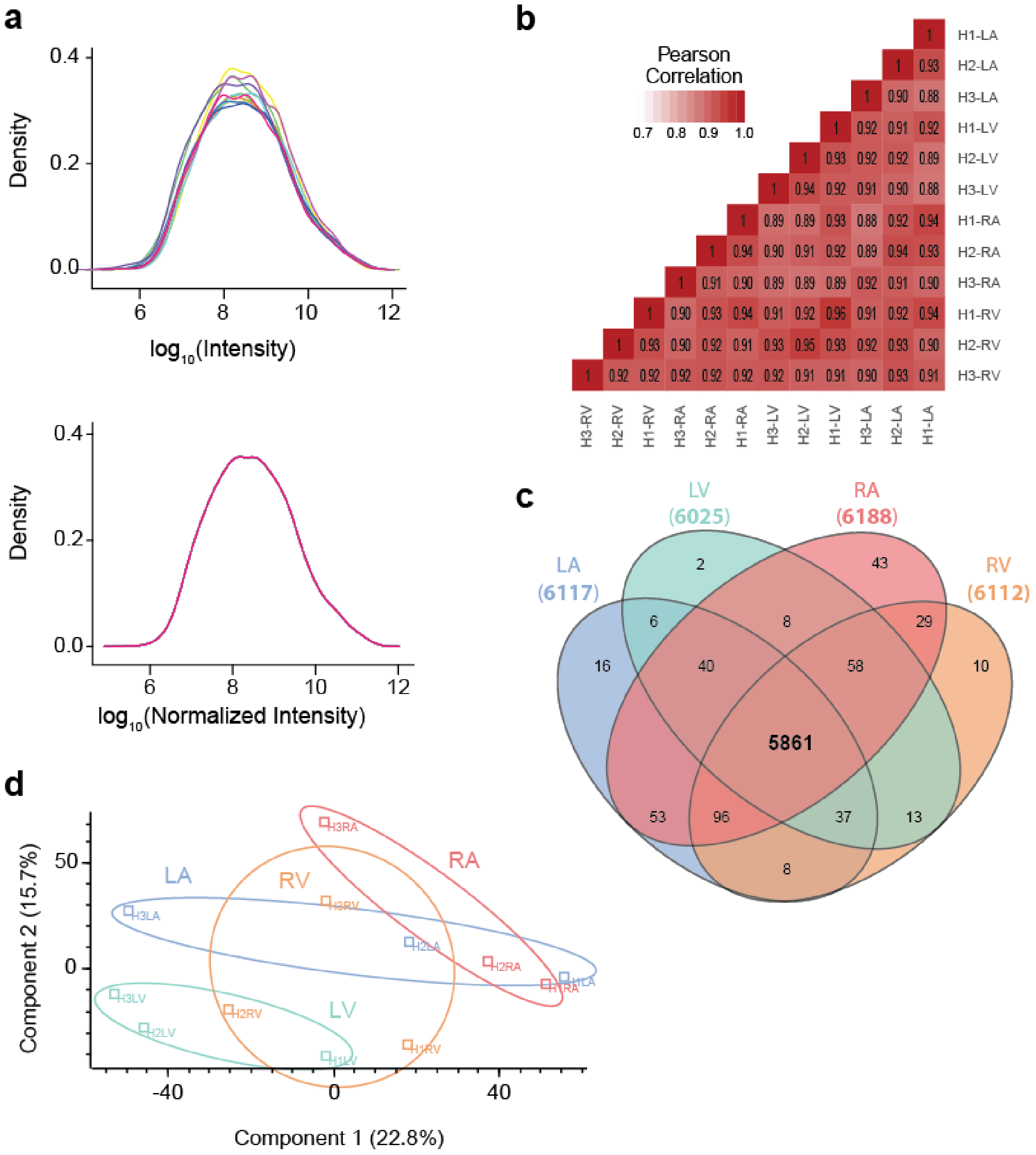

**Figure S6. Evaluation of Horse Proteome Data. a.** Density plots of intensities in log10-space displaying before (upper panel) and after quantile normalization (lower panel). **b.** Pearson correlations coefficients of log-transformed quantile normalized protein intensities across all samples. H1 through H3 denotes the three horses in the study, LA, RA, LV, and RV denotes left atrium, right atrium, left ventricle, and right ventricle respectively. **c.** Overlap of identified proteins across heart chambers are shown in a Venn diagram. More than 93% of proteins were identified in all chambers. **d**. Principal Component Analysis of samples between heart chambers or replicates in the first two principal components that explain 38.5% of the variance in the dataset.

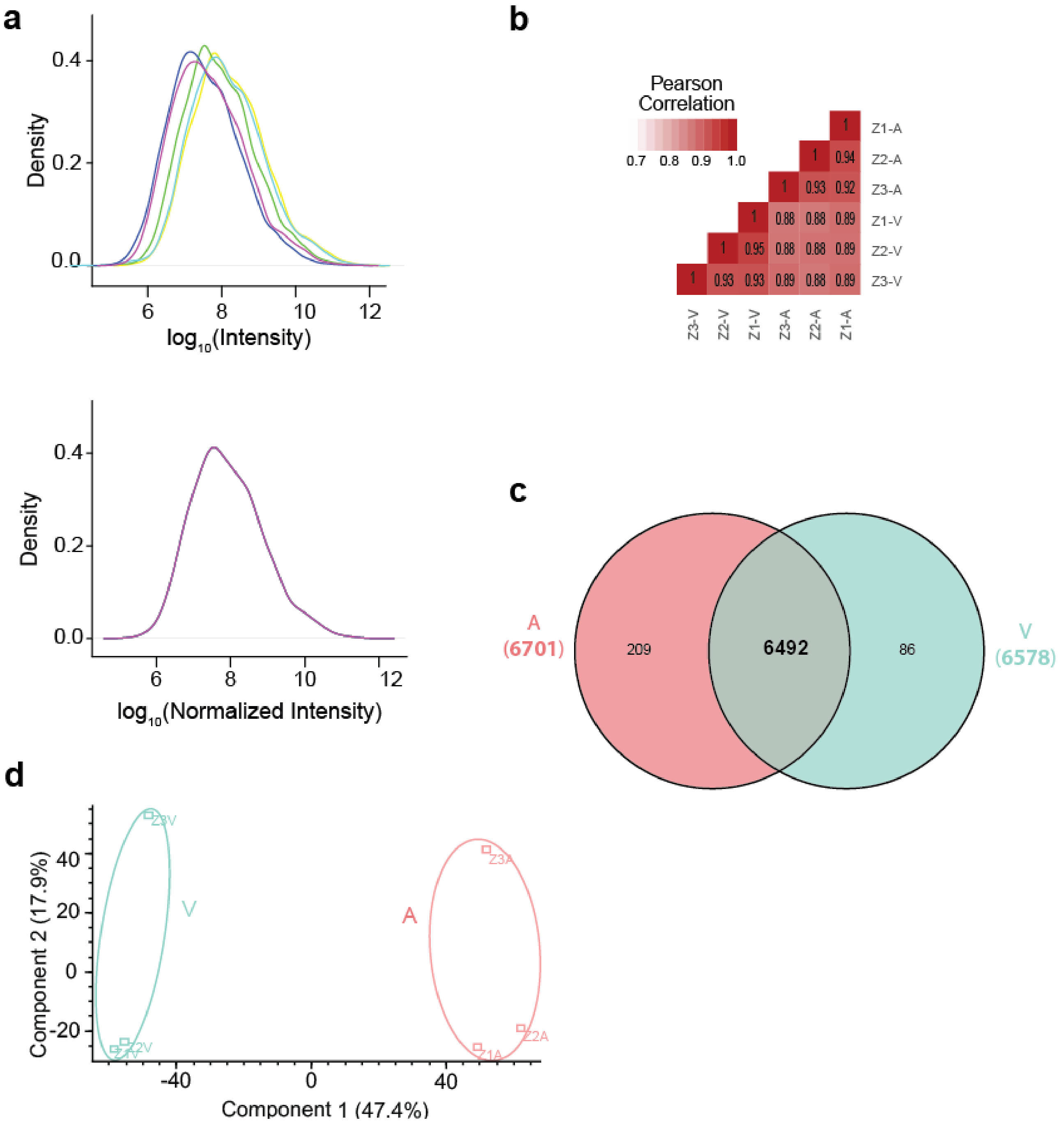

**Figure S7. Evaluation of Zebrafish Proteome Data. a.** Density plots of intensities in log10-space displayed before (upper panel) and after quantile normalization (lower panel). **b.** Pearson correlations coefficients of log-transformed quantile normalized protein intensities across all samples. Z1 through Z3 denotes the three zebrafish in the study, A and V denotes atrium and ventricle. **c.** Overlap of identified proteins across heart chambers are shown in a Venn diagram. More than 97% of proteins were identified in all chambers. **d.** Principal Component Analysis of samples shows clear distinction between atrial and ventricular samples in the first principal component that explains 47.4% of the variance in the dataset.

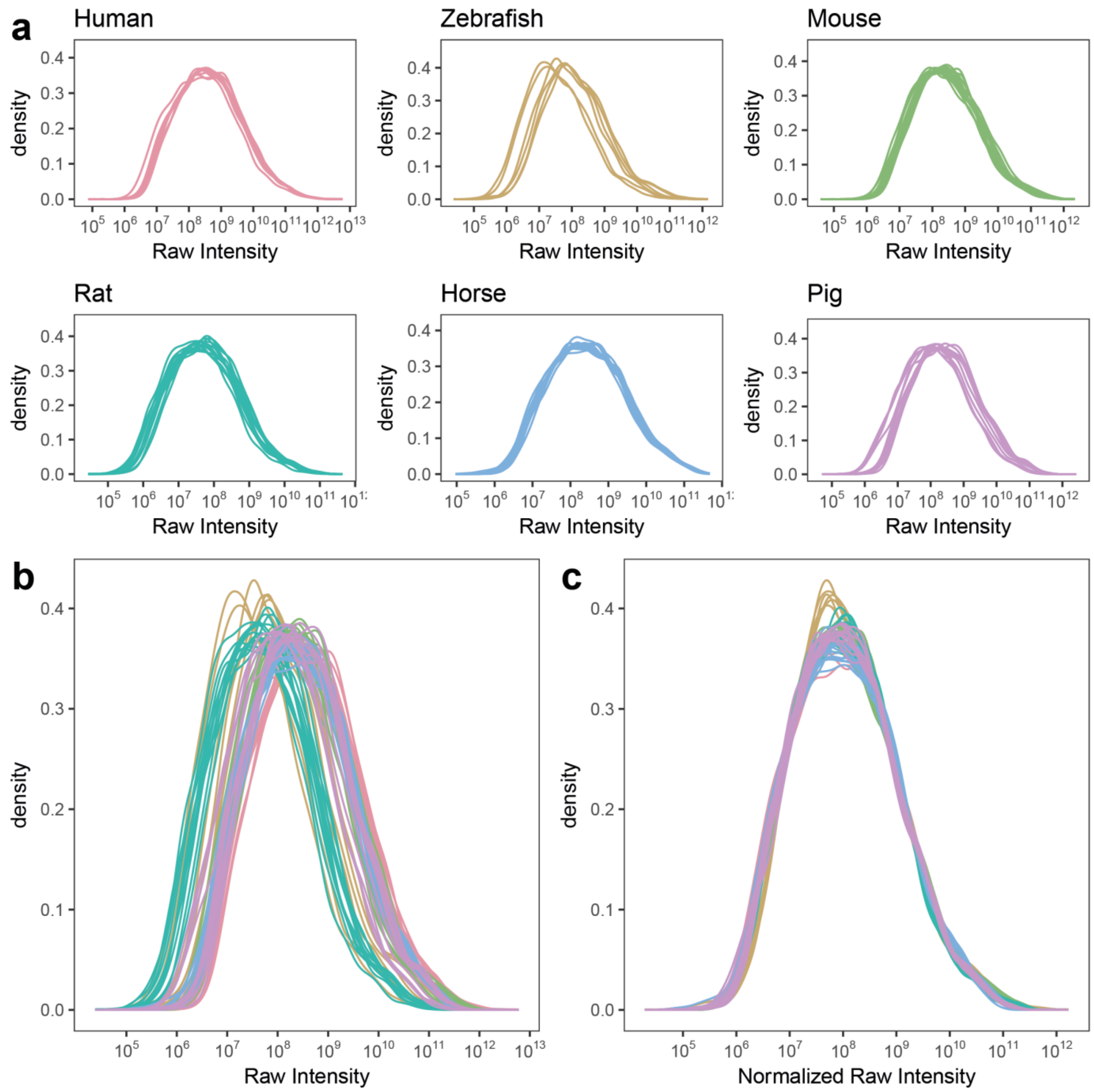

**Figure S8. Protein Intensity distributions of each species before and after data normalization. a.** Raw intensity sample distributions from each species. **b.** overlaid distributions from a. **c.** overlaid normalized raw intensity distributions by median subtraction and centering at new median 1E8 shows good overlap of data samples after normalization. This indicates sufficient similarity for comparison across species.

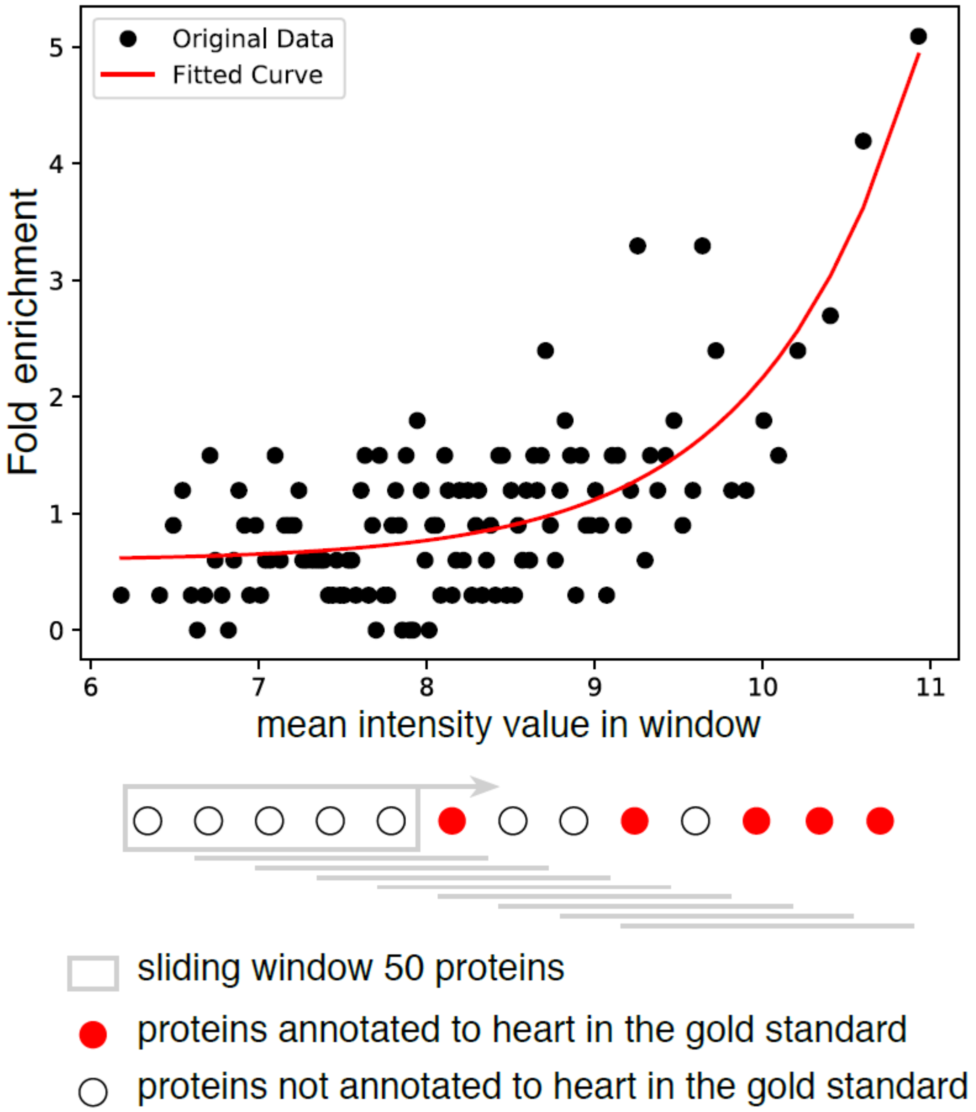

**Figure S9: Calculation of cardiac gold standard.** For representation in the database, the intensity values were translated into a multispecies confidence score by comparison to a gold standard as previously described [13]. We used the human dataset to make this comparison and calculate the new scoring scheme across species. To convert intensities into confidence scores, the agreement with the gold standard was quantified using fold enrichment. To calculate fold enrichment, we sorted proteins by intensity value and within sliding windows (window-size=50) calculated the fraction of proteins in the dataset found annotated to heart in the gold standard divided by the fraction expected when randomly sampling proteins from the gold standard. Then, we used the resulting curve (protein intensity, fold enrichment) to fit a function to translate intensities into confidence scores: – $^{-}$ , where x is the mean intensity within a sliding window of 50 proteins

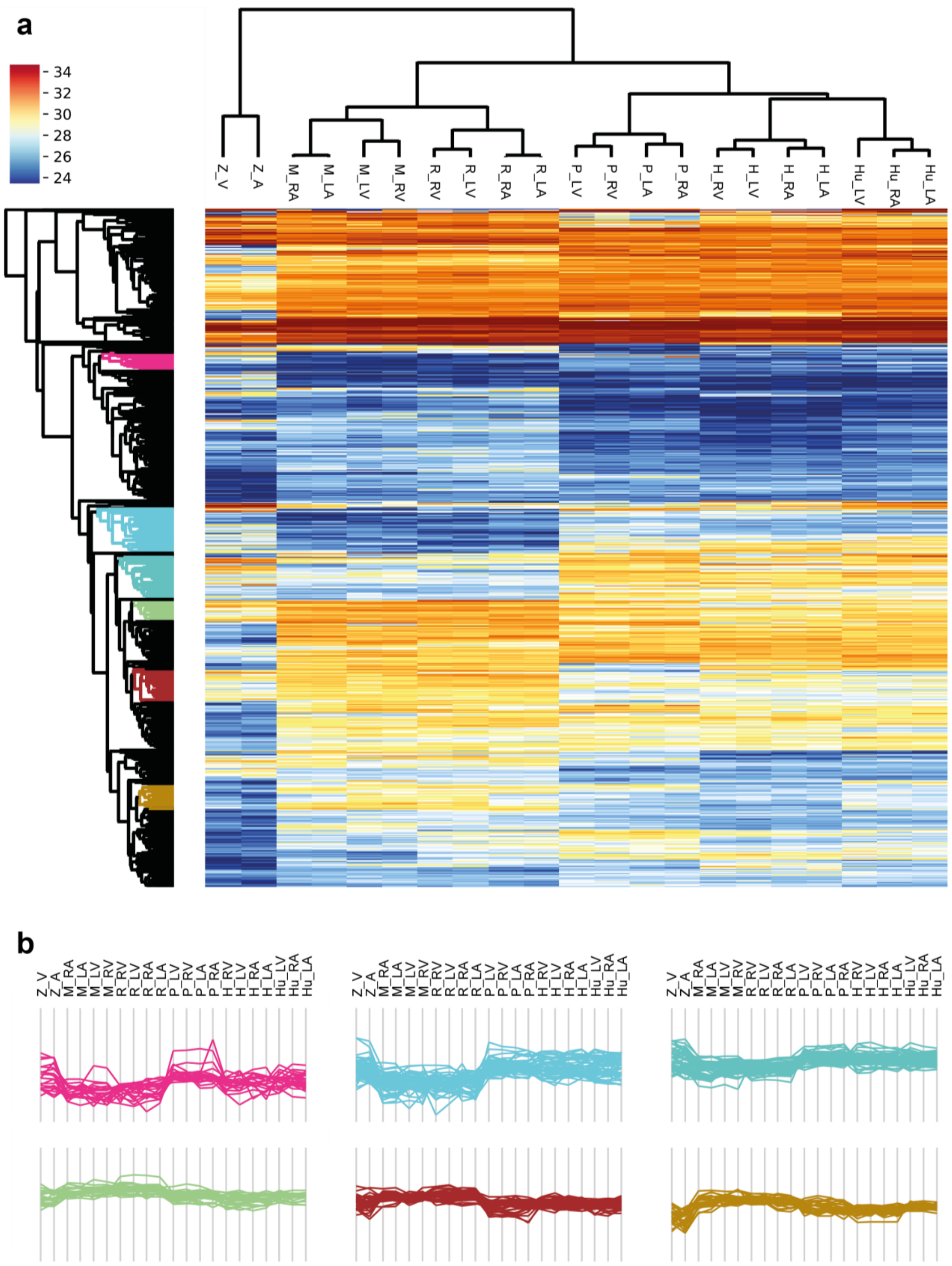

**Figure S10 Significantly differentially expressed proteins across species. a.** Subset of hierarchical clustering analysis on median protein intensities per chamber, showing all protein groups deemed significantly different based on multiple-sample ANOVA testing for differences between evolutionary groups of species (fish, small mammals, large mammals) at false discovery rate of 0.01. Six clusters were selected that showed specific up- or down-regulation of protein groups in small mammals (rat and mouse) as compared to other species. **b.** Profile plots for each cluster color-coded corresponding to panel a. Each line represents the intensity profile across samples of one orthologue group.

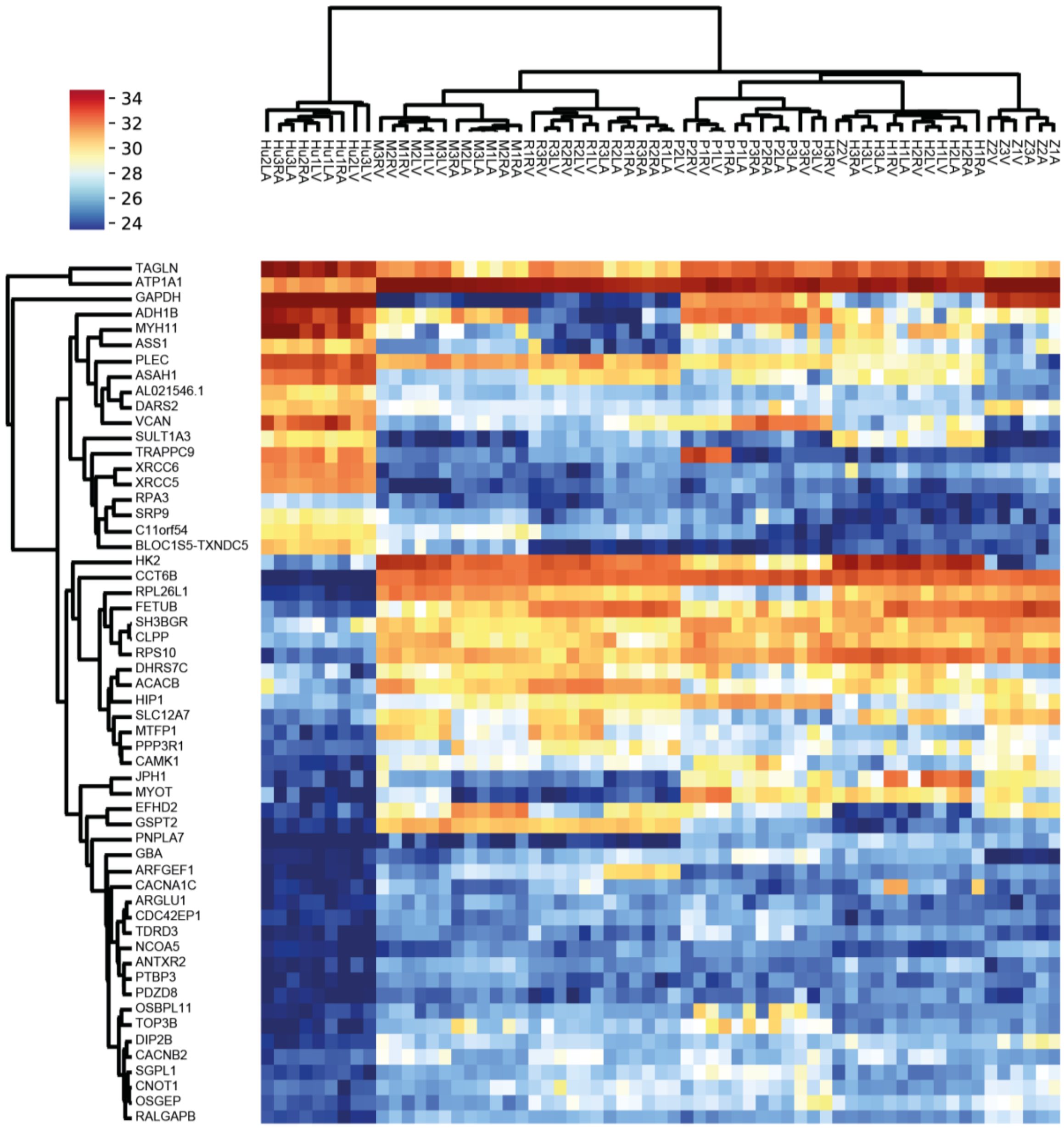

**Figure S11 Significantly different proteins in humans.** Subset of hierarchical clustering analysis on protein intensities across species, showing all protein groups deemed significantly different based on multiple-sample ANOVA testing for differences between human against all other species at false discovery rate of 0.01.

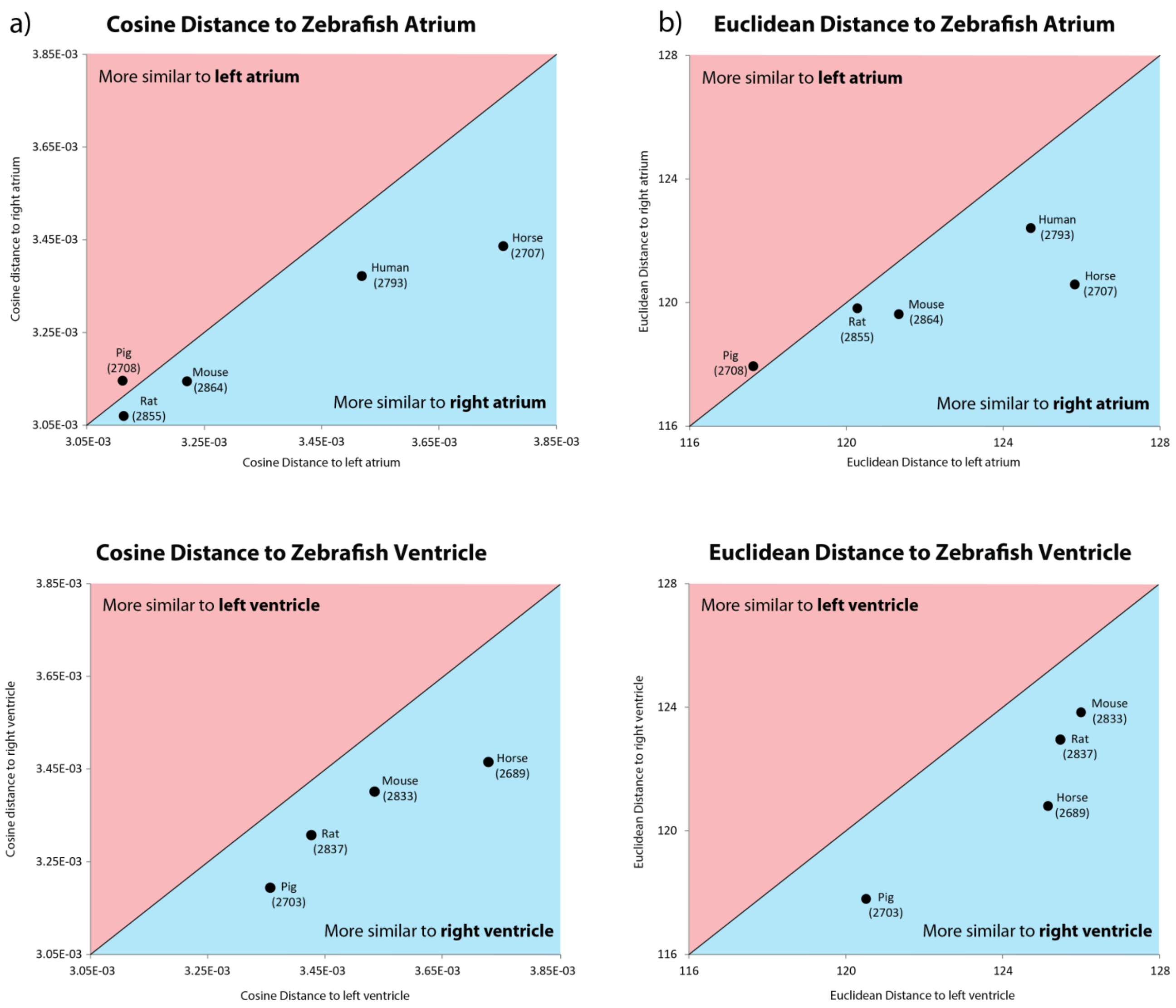

**Figure S11 Evaluation of zebra fish similarity to left/right side heart chambers**. Scatter plot of zebra fish heart chamber similarity to left/right heart chamber within mouse, rat, pig horse and human. Similarity measures cosine distance (a) and Euclidian distance (b) were used. The zebra fish atrium is more similar to the right atrium in all species except the pig, the zebra fish atrium is slightly more similar to the left pig atria by a very small margin. When comparing the zebra fish ventricle to the left/right ventricles of the mouse, rat, pig, and horse, the zebra fish displays more similarity to the right ventricle of every species.
